# A High-Throughput Physiological Functional Phenotyping System for Time- and Cost-Effective Screening of Potential Biostimulants

**DOI:** 10.1101/525592

**Authors:** Ahan Dalal, Ronny Bourstein, Nadav Haish, Itamar Shenhar, Rony Wallach, Menachem Moshelion

## Abstract

The improvement of crop productivity under abiotic stress is one of the biggest challenges faced by the agricultural scientific community. Despite extensive research, the research-to-commercial transfer rate of abiotic stress-resistant crops remains very low. This is mainly due to the complexity of genotype◻×◻environment interactions and in particular, the ability to quantify the dynamic plant physiological response profile to a dynamic environment.

Most existing phenotyping facilities collect information using robotics and automated image acquisition and analysis. However, their ability to directly measure the physiological properties of the whole plant is limited. We demonstrate a high-throughput functional phenotyping system (HFPS) that enables comparing plants’ dynamic responses to different ambient conditions in dynamic environments due to its direct and simultaneous measurement of yield-related physiological traits of plants under several treatments. The system is designed as one-to-one (1:1) plant–[sensors+controller] units, i.e., each individual plant has its own personalized sensor, controller and irrigation valves that enable (i) monitoring water-relation kinetics of each plant–environment response throughout the plant’s life cycle with high spatiotemporal resolution, (ii) a truly randomized experimental design due to multiple independent treatment scenarios for every plant, and (iii) reduction of artificial ambient perturbations due to the immobility of the plants or other objects. In addition, we propose two new resilience-quantifying-related traits that can also be phenotyped using the HFPS: transpiration recovery rate and night water reabsorption.

We use the HFPS to screen the effects of two commercial biostimulants (a seaweed extract—ICL-SW, and a metabolite formula—ICL-NewFo1) on *Capsicum annuum* under different irrigation regimes. Biostimulants are considered an alternative approach to improving crop productivity. However, their complex mode of action necessitates cost-effective pre-field phenotyping. The combination of two types of treatment (biostimulants and drought) enabled us to evaluate the precision and resolution of the system in investigating the effect of biostimulants on drought tolerance. We analyze and discuss plant behavior at different stages, and assess the penalty and trade-off between productivity and survivability. In this test case, we suggest a protocol for the screening of biostimulants’ physiological mechanisms of action.

## INTRODUCTION

To meet the food-security demands of an increasing global population, crop yields must double by 2050 (Ray et al., 2013). Despite an increase in crop productivity in the last few decades, the increased rate is not expected to match the demand, mainly due to the negative effects of climate change (abiotic environmental stresses such as drought, temperature extremes and flooding) and degrading soil quality. In fact, commercially grown crops are expected to achieve, on average, only about 50% of their potential yield under field conditions (Hatfield and Walthall, 2015; Foyer et al., 2016). In the last three decades, vast research had been invested in improving plant responses to various stresses. Nevertheless, the bench-to-field transfer rate (ratio of patents to marketed commercial seeds) of abiotic stress-resistant crops is very low, due to the high complexity of dynamic plant–environment interactions (Graff et al., 2013, Dalal et al., 2017).

### Physiological phenotyping for crop improvement

The major gap between the successful breeding and yield improvement results from the unpredictable outcome of the complex genotype ◻×◻environment interactions (Miflin 2000; Moshelion and Altman 2015). To date, the major obstacle to bridging this gap has been the lack of an efficient method for identifying and quantifying yield-related traits at early stages of plant growth across vast numbers of plants/genes (Moshelion and Altman 2015, Negin and Moshelion 2017). Another potential bottleneck is the genotype–phenotype gap. The availability of new molecular tools has enhanced the efficiency of classical breeding and crop improvement (Spindel et al., 2015; Bhat et al., 2016; Collard and Mackill 2008; Gosa et al., 2018). To achieve meaningful results in drought tolerance, molecular approaches to crop improvement must be linked to suitable phenotyping protocols at all stages, such as the screening of germplasm collections, mutant libraries, mapping populations, transgenic lines and breeding materials, and the design of OMICs and quantitative trait locus experiments (Salekdeh et al., 2009). Thus, to improve crops and to meet the challenges ahead, the genotypic view and emphasis on genomics need to be balanced by a phenocentric approach with an emphasis on phenomics, to minimize the genotype–phenotype gap (Miflin 2000). The development of a high-resolution, high-throughput diagnostic screening platform for the study of whole-plant physiological performance that serves for phenotypic screening might bridge this gap (Moshelion and Altman 2015).

Indeed, the number of phenotyping facilities has increased dramatically in the last decade. Most of these facilities collect information using robotics and automated image acquisition and analysis (White et al., 2012; Kumar et al., 2015; Ghanem et al., 2015; Fischer et al., 2014; Fiorani and Schurr 2013; Gosa et al., 2018). Nevertheless, the quest for more detailed and in-depth phenotyping of the dynamic genotype◻×◻environment interactions and plant stress responses (in particular during drought) has put the capability of the existing methods into question (Ghanem et al., 2015; Li et al., 2014; Halperin et al., 2017; Rahaman et al., 2015; reviewed by Gosa et al., 2018). Herein, we demonstrate a high-throughput functional phenotyping system (HFPS) composed of gravimetric systems that enable us to compare plants’ dynamic responses to different ambient conditions in dynamic environments, due to its direct and simultaneous measurement of the yield-related physiological traits of all plants under several treatments.

### Phenotyping for biostimulants in drought response

Apart from the traditional strategies to improve crop productivity under an uncertain environment and abiotic stress, an alternative approach is evolving. This approach considers the use of organic molecules, externally applied to the plant at low concentrations, to stimulate many aspects of growth and development, pathogen defense, stress tolerance and reproductive development. These organic molecules, collectively termed biostimulants, have become more and more common in the global market in the last two and a half decades (reviewed by Yakhin et al., 2017). Biostimulants have been defined in many different ways. In the scientific literature, the term biostimulant was first defined by Kauffman et al. (2007) in a peer-reviewed paper, with modifications: “biostimulants are materials, other than fertilizers, that promote plant growth when applied in low quantities” (reviewed by du Jardin, 2015). However, the definition of biostimulants adopted by the European Biostimulants Industry Council specifies that these materials should not function by virtue of the presence of essential mineral elements, known plant hormones or disease-suppressive molecules (Brown and Saa 2015). Recently, biostimulants were defined by Yakhin et al. (2017) as “a formulated product of biological origin that improves plant productivity as a consequence of the novel, or emergent properties of the complex of constituents, and not as a sole consequence of the presence of known essential plant nutrients, plant growth regulators, or plant protective compounds.” However, due to their complex composition and diversity, biostimulants are classified differently by different research groups. Many categorize biostimulants based on the natural raw materials used, the origin of their active ingredients and modes of action, inclusion or exclusion of microorganisms, and/or mode of action of the biostimulant (Ikrina and Kolbin 2004; Basak 2008; Du Jardin 2012; Bulgari et al. 2015; Yakhin et al., 2017).

Biostimulants are used in all stages of agriculture, namely, in seed treatments, during plant growth, and postharvest. They are applied both as foliar sprays and through the soil. Biostimulants may function in various ways, as comprehensively summarized by Yakhin et al. (2017). Their mechanism of action may comprise activation of nitrogen metabolism or phosphorus release from soils, generic stimulation of soil microbial activity, or stimulation of root growth and enhanced plant establishment. They stimulate plant growth by enhancing plant metabolism, stimulating germination, improving photosynthesis, and/or increasing the absorption of nutrients from the soil, thus increasing plant productivity. Studies have shown a clear protective role of a diverse number of biostimulants against abiotic stress, as reviewed by Van Oosten et al. (2017). Nevertheless, and despite the extensive literature suggesting that biostimulants decrease the effects of abiotic stress (and in particular drought stress), information regarding their physiological mechanisms of action is limited. The large number of potential candidate biostimulants and the need to elucidate their particular modes of action, optimal concentrations, and types of application, create a substantial bottleneck in the research and development of new biostimulant products. High-throughput phenotyping technologies have been successfully employed in some aspects of plant breeding (Araus and Cairns, 2014; Tardieu et al., 2017), but their application to assess plant biostimulant action has been limited (Petrozza et al., 2014; reviewed by Rouphael et al., 2018), despite the potential benefits of using these technologies in biostimulant product screening (Rouphael et al., 2018).

In this study, we tested the effectiveness of physiological phenotyping for understanding the physiological ‘mode of action’ of biostimulant activity on the whole plant’s drought response. We tested the impact of biostimulants on several quantitative yield-related physiological traits: transpiration rate, growth rate, and water-use efficiency (WUE).

## MATERIALS AND METHODS

### Plant Material

The seeds of pepper (*Capsicum annuum* var. Rita) were obtained from Zeraim Gedera-Syngenta, Israel. For germination, the seeds were sown in a tray with 10-mL cones filled with commercial growing medium (Matza Gan, Shaham, Givat-Ada, Israel), composed of (w/w) 55% peat, 20% tuff and 25% puffed coconut coir fiber. The trays were well irrigated and kept in the same greenhouse (on side-tables) where the experiment was performed. When the seedlings were 4 weeks old, the growing medium was carefully washed off (to avoid root damage) the seedling roots and the seedlings were immediately transferred to 4-L pots filled with 20/30 sand (Negev Industrial Minerals Ltd., Israel). The numbers 20/30 refer to the upper and lower size of the mesh screen through which the sand was passed (20 = 20 squares across one linear inch of screen), resulting in a sand particle size of between 0.595 and 0.841 mm. The volumetric water content (VWC) of the freely drained substrate, noted as pot capacity, was ~24% (for details, see Experimental Setup section).

### The Physiological Phenotyping Platform

The experiment was conducted in June–July 2018 in a commercial-like greenhouse located at the Faculty of Agriculture, Food and Environment in Rehovot, Israel. The greenhouse temperature was controlled using fans that blow air through a moist mattress, keeping it below 38°C. The temperature and relative humidity (RH) were 21–38°C and 30–80%, respectively. The plants were grown under natural light (midday maximum of 1300 μmol◻ s^-1^ m^-2^), representative values for natural conditions during the summer in the central part of Israel, including Rehovot. The temperature, RH, photosynthetically active radiation, barometric pressure and vapor pressure deficit in the greenhouse were continuously monitored by Plantarray meteorological station (Plant-Ditech Ltd., Israel).

The functional phenotyping system Plantarray 3.0 platform (Plant-Ditech) was used to monitor the plants’ performance during the entire experimental period by controlling the schedule and quantity of irrigation. This platform (Figure 1A and Supplementary Figure 1), which enables performing high-throughput physiological functional phenotyping, includes 72 units of highly sensitive, temperature-compensated load cells that are used as weighing lysimeters. Each unit is connected to its personalized controller, which collects the data and controls the irrigation to each plant separately. An independent controller for each pot enables tight feedback irrigation, based on the plant’s transpiration rate. Each controller unit is connected to its neighbor for serial data collection and loading to a server. A pot with a single plant is placed on each load cell (for more details, see Experimental Setup section). The data were analyzed by SPAC-analytics (Plant-Ditech), a designated online web-based software that enables viewing and analyzing the real-time data collected from the Plantarray system.

**FIGURE 1.**
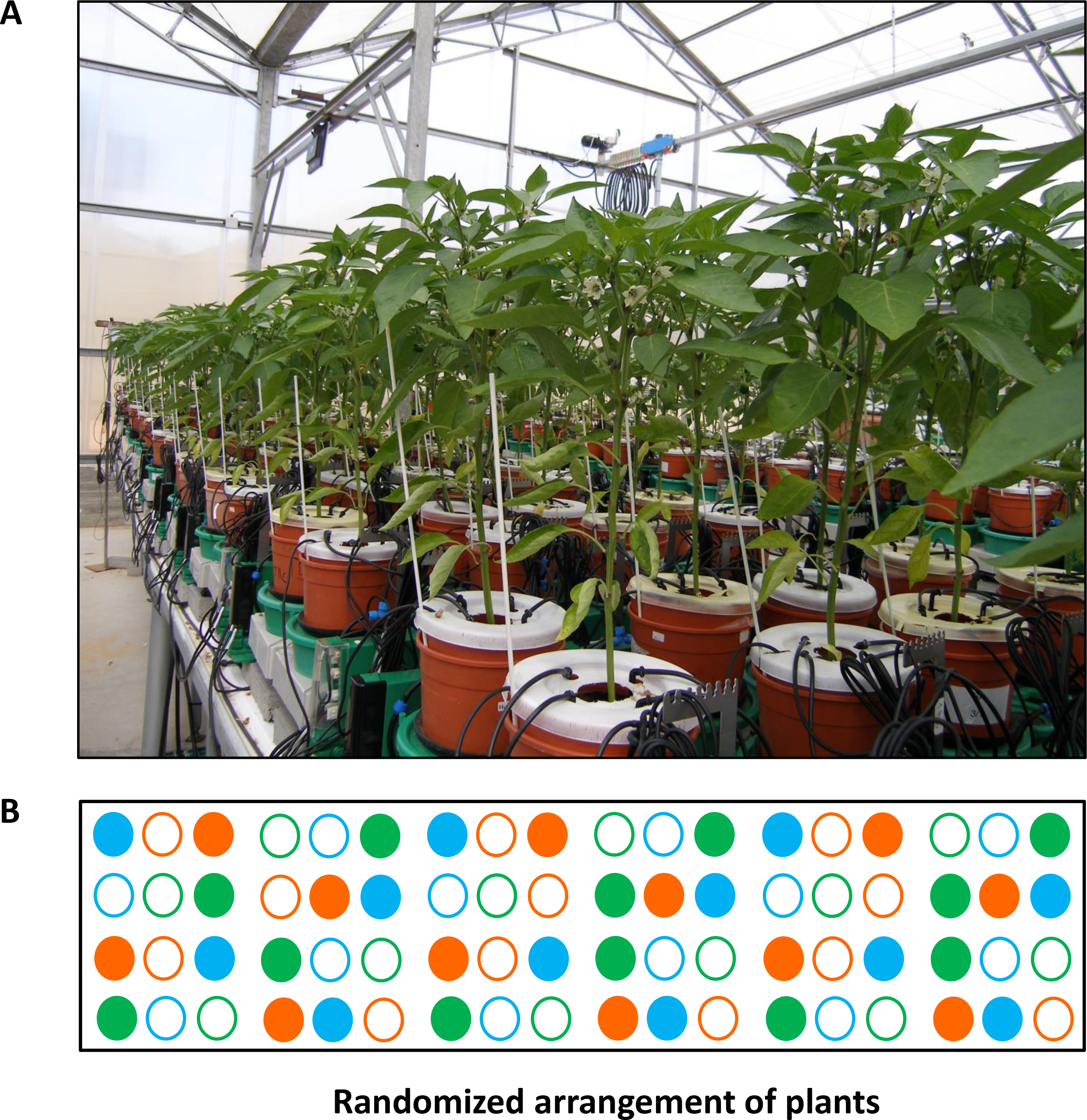
Experimental setup. **(A)** View of the randomized experimental setup array consisting of 72 measuring units loaded with *Capsicum annuum*. **(B)** Block diagram of the system. Solid circles – well-irrigated plants; empty circles – plants subjected to the drought-recovery scenario. Green – ICL-SW-treated plants, orange – ICL-NewFo1-treated plants, blue – control (no biostimulants) plants. Note that all pot surfaces were covered to reduce evaporation, and irrigation was injected into the soil via multi-outlet drippers to ensure even distribution of fertigation and biostimulants (see Supplementary Figure 1).

### Nutrition and Treatments

The composition of the nutrients supplied to the plants by the irrigation system (fertigation) is provided in Table 1. Two different commercial biostimulants were used: seaweed extract (ICL-SW) and a metabolite extract formula (ICL-NewFo1) (both supplied and produced by ICL Specialty Fertilizers, Holland). The biostimulants were prepared in two different containers that were placed on two additional load cells to precisely track their application. The biostimulants were provided to the plants together with the nutrients via the controlled irrigation system (Supplementary Figure 1). The biostimulant concentration and dosage were as per the manufacturer’s instructions: ICL-NewFo1 (3.53 mg/L) was provided daily and ICL-SW (0.133 mg/L) once a week.

The experiment lasted 36 days and included two treatments: (i) ample irrigation that aimed to provide non-stressed conditions for the plants throughout the experiment (termed well-irrigated plants), (ii) controlled drought (days 13–30) preceded by a period of ample irrigation, noted as pretreatment (days 1–12), and followed by resumption of ample irrigation (recovery period) (see Figure 2B and Experimental Setup section for details). The treatments included ICL-SW, ICL-NewFo1 or no biostimulants (control). Overall, we had six different experimental groups: three with ample irrigation (control–well irrigated, ICL-SW–well irrigated and ICL-NewFo1– well irrigated) and three groups subjected to drought (control–drought, ICL-SW– drought, and ICL-NewFo1–drought). Each of these groups consisted of 8–12 repetitions (plants) that were arranged in a randomized fashion on the array to ensure uniform exposure of all groups, thereby overcoming the inherent variations in ambient conditions (Figure 1B).

**FIGURE 2.**
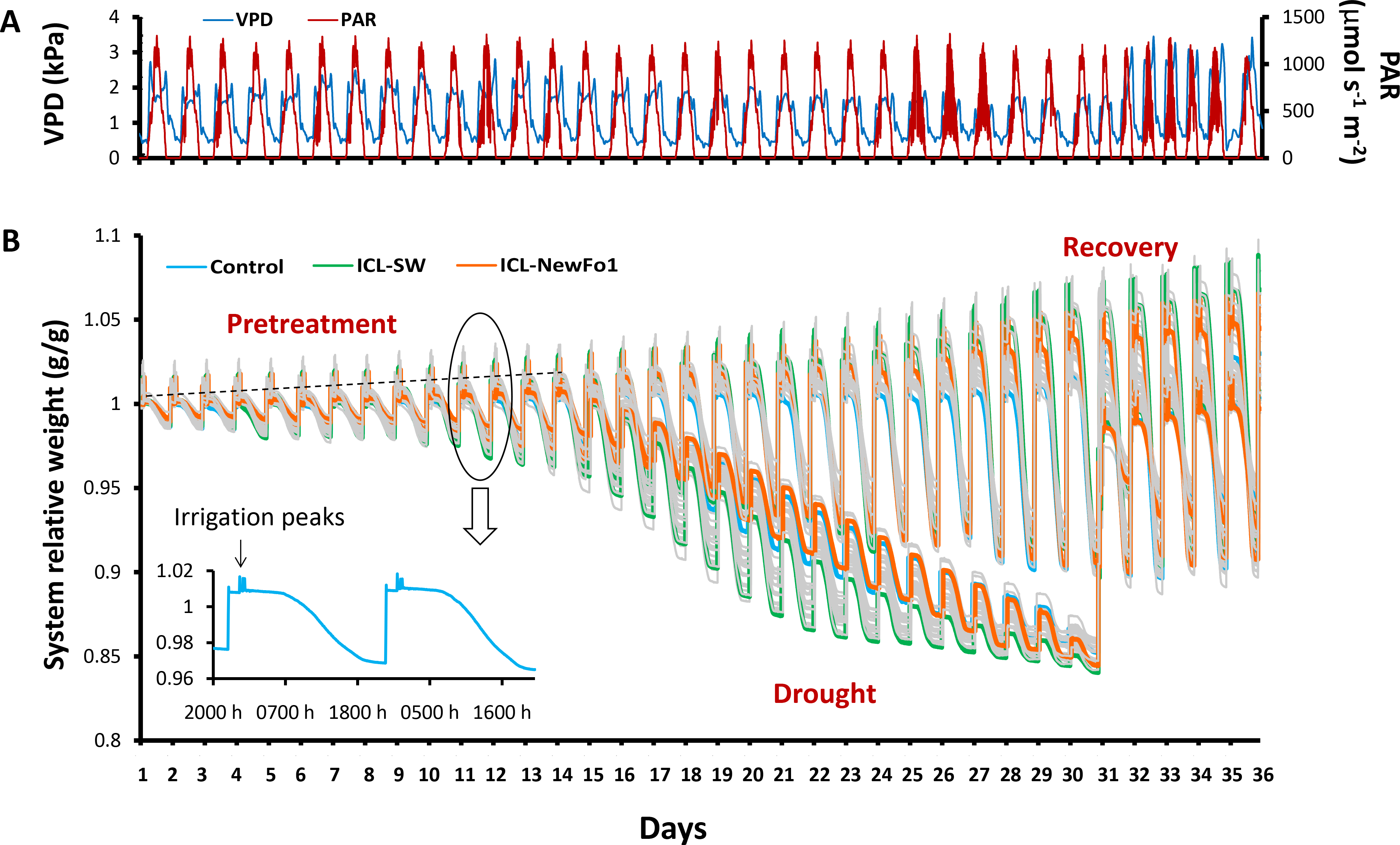
Atmospheric conditions and experimental progress represented as system relative weight throughout the experiment. **(A)** Daily vapor pressure deficit (VPD) and photosynthetically active radiation (PAR) during 36 consecutive days of experiment. **(B)** Raw data showing variation in the weight of the plants (relative to their respective initial weight) over the course of the experiment. Each line represents one plant/pot. During the day, the plant transpires and therefore the system loses weight, seen as a slope in the line curves. The pots were irrigated four times per night (each time to pot capacity), seen as peaks in the line curves. The irrigation was followed by drainage to reach water saturation (nighttime baseline). Note that there is no weight loss during the night. The increase in the nighttime baseline (dashed line) every day results from an increase in plant biomass. During pretreatment, all of the plants were well irrigated; from day 13, half of the plants were exposed to differential drought to reach a similar degree of stress, while the other half continued to be well-irrigated till the end of the experiment. On day 31, the water-deprived plants were recovered and continued to be well-irrigated till the end of the experiment. The three colored lines represent a single plant from each of the three groups: blue line – untreated (with biostimulants) control plants; green line – ICL-SW-treated plants; orange line – ICL-NewFo1-treated plants. Note the different drought-response behaviors of the different plants. The inset figure presents system relative weight change of one plant/pot for two consecutive days.

### Experimental Setup

The experimental setup was generally similar to Halperin et al. (2017) with some modifications. Briefly, before the start of the experiment, all load-cell units were calibrated for reading accuracy and drift level under constant load weights (1 kg and 5 kg). Sand was used as the growing substrate because (i) it is an inert substance (sand is free of any nutrients, helping to precisely understand the effect of any chemical applied externally through irrigation), (ii) it is easily washed off the roots (helping to study the roots after the completion of the experiment), and (iii) pot capacity is reached rapidly with a repeatable pattern after each irrigation (at the end of free drainage), helping to study the plants’ short-term resilience trait, noted as “night water reabsorption” (see Measurement of Quantitative Physiological Traits section for details). The sand in all of the pots was washed thoroughly several times prior to transfer of the seedlings. Each pot was placed into a Plantarray plastic drainage container on a lysimeter. The container fit the pot size to prevent evaporation. The container has orifices on its side walls at different heights to enable different water levels after drainage of excess water following irrigation. Evaporation from the sand surface was prevented by a plastic cover with a circle cut out at its center through which the plants grew.

Each pot was irrigated by multi-outlet dripper assemblies (Netafim, Israel) that were pushed into the soil to ensure that the medium in the pot was uniformly wetted at the end of the free drainage period following each irrigation event. Irrigations were programmed to run during the night in four consecutive pulses. A 2-h interval was maintained between the first irrigation pulse and the last three. This irrigation regime enabled determining the plants’ night water reabsorption, one of the traits indicating plant resilience (see Measurement of Quantitative Physiological Traits section). The amount of water left in the drainage containers underneath the pots at the end of the irrigation events was intended to provide water to the well-irrigated plants beyond the water volume at pot capacity. The associated monotonic weight decrease throughout the day hours was essential for the calculation of the different physiological traits by the data-analysis algorithms.

The drought treatment started on day 13 and ended when the plants’ daily transpiration had reached ~80 mL per day. To prevent rapid depletion of the water in the sandy soil in the pots, we conducted gradual deficit irrigation that reduces the irrigation levels every day to 80% of the previous day’s transpiration, for each plant separately and independently (using Plantarray’s automated feedback irrigation system; Figure 2B).

### Measurement of Quantitative Physiological Traits

The following water-relations kinetics and quantitative physiological traits of the plants were determined simultaneously, following Halperin et al.’s (2017) protocols and equations implemented in the SPAC-analytics software: daily transpiration, transpiration rate, normalized transpiration (E), transpiration rate vs. calculated VWC using a piecewise linear fit, and WUE. Cumulative daily transpiration was calculated as the sum of daily transpiration for all 36 days of the experiment. The VWC in the sand medium was calculated by a mass balance between the system weight at pot capacity when free drainage ceases and its concurrent weight.

The estimated plant weight at the beginning of the experiment was calculated as the difference between the total system weight and the sum of the tare weight of pot + drainage container, weight of soil at pot capacity, and weight of water in the drainage container at the end of the free drainage. The plant weight at the end of a growth period (calculated plant weight) was calculated as the sum of the initial plant weight and the multiplication of the cumulative transpiration during the period by the WUE. The latter, determined as the ratio between the daily weight gain and the daily transpiration during that day, was calculated automatically on a daily basis by the SPAC-analytics software. Note that the WUE approached a constant value during the pretreatment period.

The plant’s recovery from drought was described by the rate at which the plant gained weight following resumption of irrigation (recovery stage). The physiological trait representing the plant’s transpiration recovery from drought was determined as the ratio between the slope of the daily transpiration increase during the recovery phase (recovery slope) and the slope of the daily transpiration decrease during the drought period (stress degree). The slopes were calculated using a linear regression.

The night water reabsorption trait was determined as the difference in system weight between the end of the last and first irrigations of a given irrigation event (i.e. single night), representing the water absorbed by the plant during the very short period when transpiration is practically negligible. This calculation is based on the fact that the drainage of surplus water in sand is rapid and pot capacity is reached prior to the subsequent irrigation (Supplementary Figure 2). We considered the plants’ short-term water reabsorption capability during the recovery stage to be an additional physiological trait representing the plant’s resilience to drought. Note that the water reabsorption by the plant during the night hours was normalized to its weight.

The recovery stage lasted 6 days, after which the experiment was stopped. As pepper is an indeterminate plant, it did not reach its full yield capacity. Consequently, the experiment was terminated at this stage as the treatment conducted to that point had a direct effect on the existing fruit. The shoots and fruit were harvested from ~10-week-old plants, irrespective of their size, in the early morning hours. The fresh shoot weight was calculated by the system as the difference in actual gravimetric weight between the day of shoot harvest at 0400 h (at the end of the last irrigation) and the following day at the same time. The fruit were collected from the harvested shoot and counted. The fruit and shoots (without fruit) were weighed when a constant weight had been reached during drying in a hot air oven at 60°C. The roots were collected from the pots, washed thoroughly to remove the sand particles, and dried in a hot air oven at 60°C until no further reduction in weight was measured, and finally weighed. The total dry plant weight is the sum of dry shoot weight, dry root weight and dry fruit weight.

### Statistical Analysis

Means were compared using analysis of variance (ANOVA) and Student’s *t*-test (noted in the figure legends) in JMP Pro 14 software.

## RESULTS

A randomized experimental design was performed to quantitatively compare the impacts of two biostimulants (seaweed extract ICL-SW and metabolite formula ICL-NewFo1) on the plant’s key physiological traits. The effects of the two biostimulants were compared to controls (no biostimulant) under two irrigation scenarios: (i) well irrigated, and (ii) drought stress starting with a well-irrigated period, then a controlled drought phase and a successive recovery period (Figure 2B).

### Biostimulants Affect Plant Water Loss

Daily transpiration increased gradually for all six groups during the well-irrigated period (pretreatment; Figure 3A). Conversely, daily transpiration and VWC in the pot gradually decreased throughout the drought period that started on day 13 of the experiment (Figures 3A and B, respectively). Daily transpiration and VWC and increased sharply upon irrigation resumption on day 31 of the experiment (recovery period) (Figures 3A and B, respectively). The physiological drought point (defined as the soil VWC value that begins to limit transpiration rate [critical VWC, θ_critical_ (θc)]) was determined for the plants subjected to drought (Figure 3C). A θc = 0.15 was obtained for the control and two biostimulant treatments, but due to the different pattern of VWC decrease in the ICL-SW-treated plants compared to the other two groups (Figure 3B), they reached θc on different days. The θc for the control and ICL-NewFo1-treated plants was reached on day 22.5, and on day 21 for the ICL-SW-treated plants (Figure 3B,C). The impact of drought on the daily transpiration rate pattern of the treated and untreated plants relative to that of the three well-irrigated groups is illustrated in Figure 3D for days 27–29, revealing that the ICL-SW-treated plants experienced a significantly lower midday (between 1200 and 1400 h) transpiration rate under drought but reached a significantly higher transpiration rate under full irrigation (Figure 3E). Under ample irrigation, the ICL-NewFo1-treated plants had a significantly higher transpiration rate than the control plants, and a similar reduction in transpiration rate under drought (Figure 3E).

**FIGURE 3.**
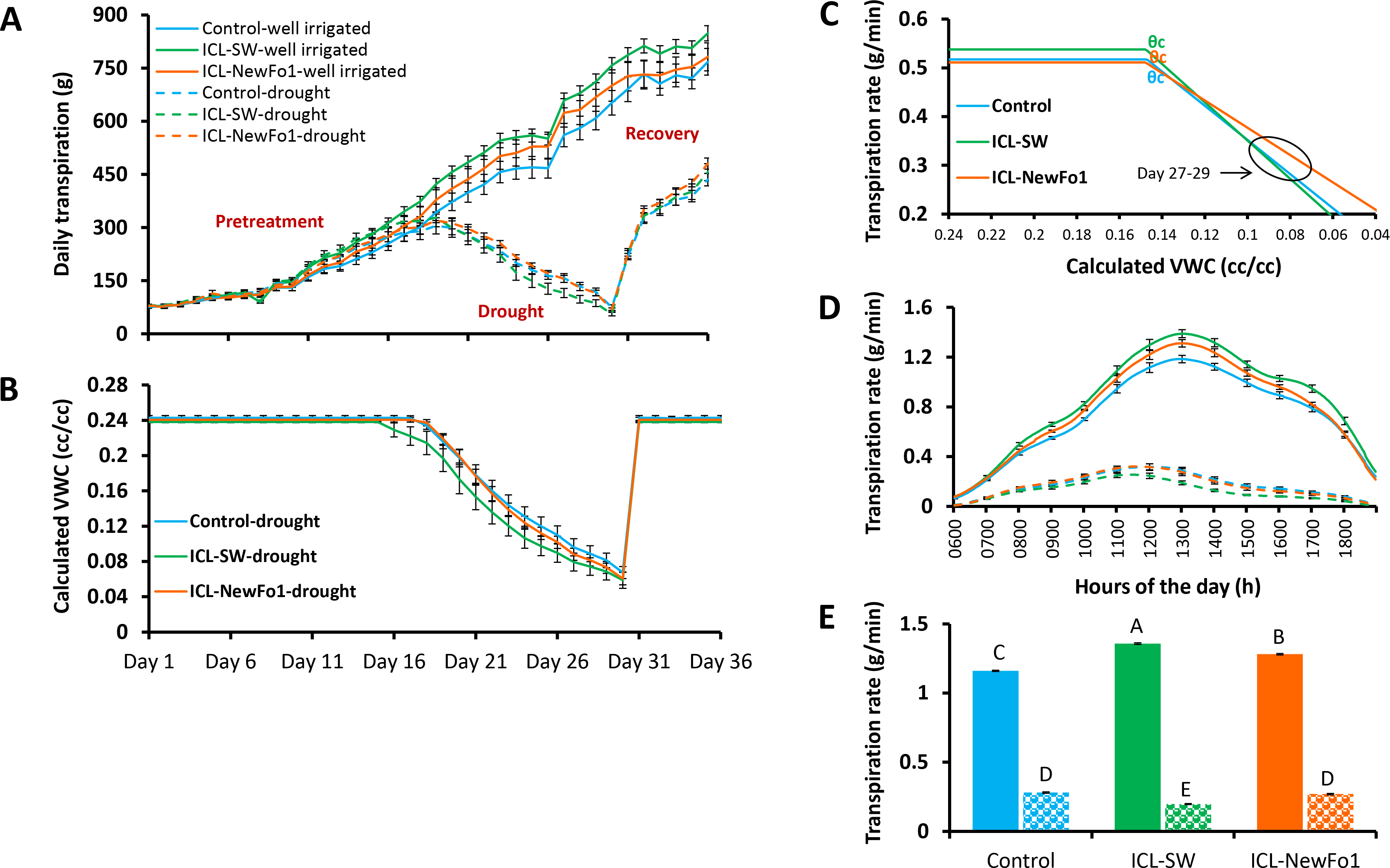
Effect of biostimulants on plant transpiration. **(A)** Mean ± SE continuous daily whole-plant transpiration during the entire experimental period (36 days). **(B)** Mean ± SE calculated volumetric water content (VWC) of the water-deprived plants throughout the experiment. **(C)** Piecewise linear fit between transpiration rate and calculated VWC for the plants subjected to drought treatment. **(D)** Mean ± SE diurnal transpiration rate from 0600 to 1900 h during the late drought phase (day 27–29). **(E)** Mean ± SE transpiration rate for days 27–29 from 1200 to 1400 h. Blue bars – no biostimulant control plants; green bars – ICL-SW-treated plants; orange bars – ICL-NewFo1-treated plants. Solid bars – well-irrigated conditions; stippled bars – drought conditions. Groups were compared using ANOVA by Tukey HSD test. Different letters above columns represent significant differences (*P* < 0.05). Each mean ± SE is from at least 8 plants per group.

### Biostimulants Enhance Biomass and WUE

Transpiration was normalized to biomass by using the calculated plant weight for the entire experimental period (36 days) for all six groups (Figure 4A). The rate of plant weight gain during the well-irrigated period (pretreatment) was similar for all six groups, and decreased during the drought period for the three drought-stressed groups. The rate of weight gain for the latter groups began to increase again during the recovery period (Figure 4A). Nevertheless, the higher rate of weight gain for the ICL-SW-treated plants during this latter period resulted in significantly higher dry shoot biomass than for controls at the end of the experiment, probably due to the cumulative effect of this trend (Figure 4B). The correlation between shoot dry biomass and cumulative daily transpiration, which is, in fact, the dry-weight-related WUE, was relatively high (R^2^ > 0.8) for both the well-irrigated and water-deprived plants (Figure 4C). Plant transpiration normalized to plant weight, E (Figure 4D), was low for biostimulant-treated plants under both well-irrigated and drought conditions compared to its value for untreated controls. Here again, the ICL-SW-treated plants showed significantly lowest midday E under drought (in accordance with the transpiration rate, Figure 3E). The higher measured transpiration rates (Figure 3E) and higher dry shoot biomass (Figure 4B) for the biostimulant-treated plants compared to the controls under ample irrigation indicate an improvement in fresh weight-related WUE. However, this improvement (increase of ~18% for ICL-SW-treated and ~14% for ICL-NewFo1-treated plants) was not significant (*P*-value for ICL-SW was 0.067 and for ICL-NewFo1, 0.16; Figure 4E).

**FIGURE 4.**
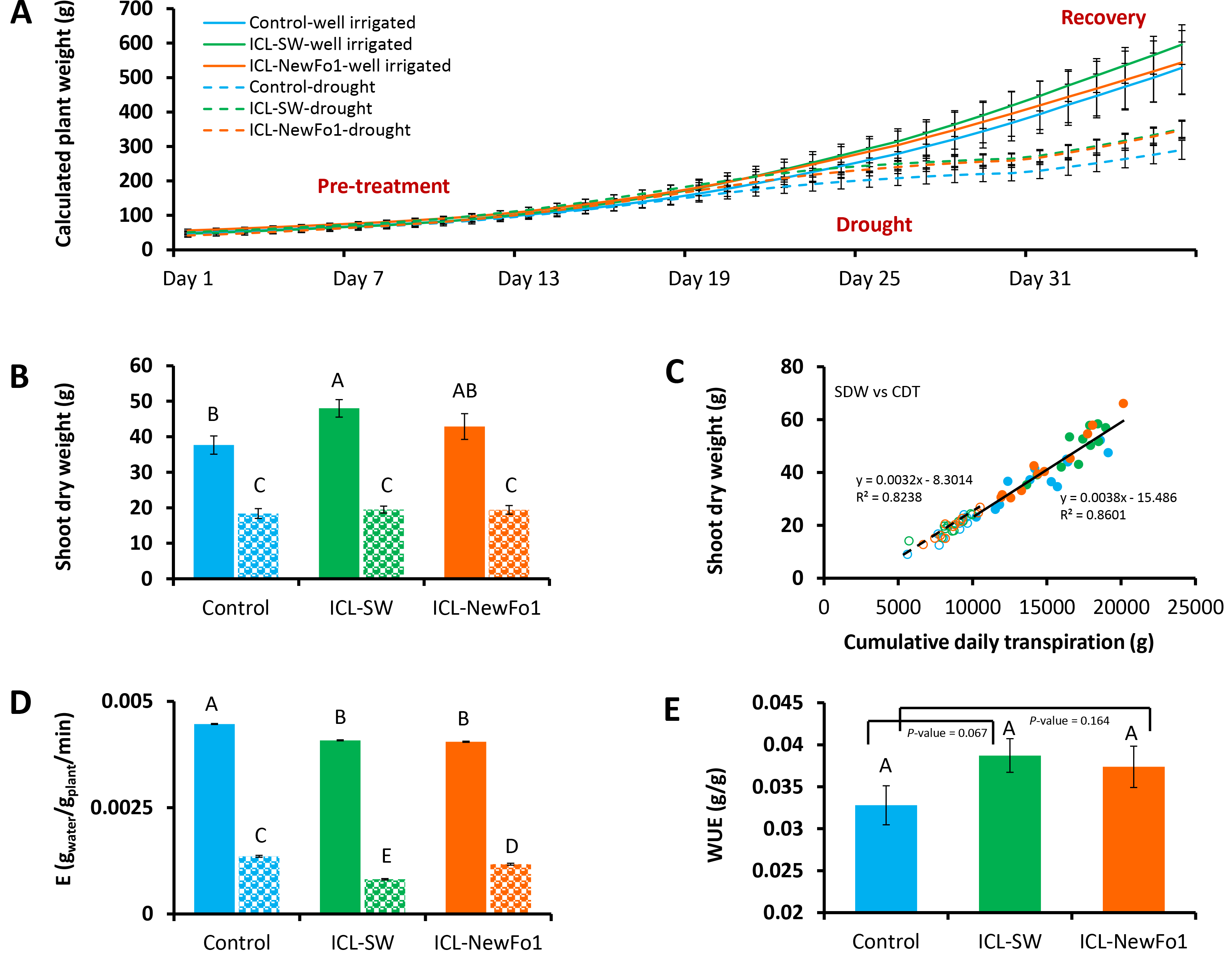
**(A)** Mean ± SE calculated whole-plant weight during the entire experimental period. **(B)** Mean ± SE shoot dry weight, harvested at the end of the experiment. **(C)** Correlation between shoot dry weight and cumulative daily transpiration. **(D)** Midday mean ± SE E (transpiration rate normalized to plant biomass) for days 27–29 from 1200 to 1400 h. **(E)** Mean ± SE water-use efficiency (WUE). Blue bars – untreated (with biostimulants) control plants; green bars – ICL-SW-treated plants; orange bars – ICL-NewFo1-treated plants. Solid bars – well-irrigated conditions; stippled bars – drought conditions. Groups were compared using ANOVA by Tukey HSD test and Student’s *t*◻test. Different letters above columns represent significant differences (*P* < 0.05). Each mean ± SE is from at least 8 plants per group.

### Biostimulant Effect on Plant Resilience

The two considered traits for an estimation of the plants’ recovery from drought stress, i.e., resilience, were: (i) whole-plant transpiration recovery: the rate at which the daily transpiration increases following irrigation resumption was compared to the rate at which the daily transpiration decreases during the drought period. For the sake of simplicity, both rates were determined as the linear regression of the respective data points (Figure 5A), showing that ICL-SW reduced plant resilience compared to control plants (Figure 5B); (ii) the night water reabsorption (namely, regaining the water that was lost during the day; see Supplementary Figure 2) for the pretreatment and recovery periods, depicted in Figures 5C,D and 5E,F, respectively. The night water reabsorption during the pretreatment period was significantly higher for the biostimulant-treated plants compared to the control, with the highest values for the ICL-NewFo1-treated plants (Figure 5C). The drought stress reduced night water reabsorption capability during recovery for all three groups. Nevertheless, compared to the control, the biostimulants improved the reabsorption capability during recovery, with significantly highest capability for ICL-NewFo1-treated plants (Figure 5E). A similar trend was observed when the night water reabsorption was normalized to the plant weight, with significantly highest reabsorption capability in ICL-NewFo1-treated plants compared to the control (Figures 5D,F).

**FIGURE 5.**
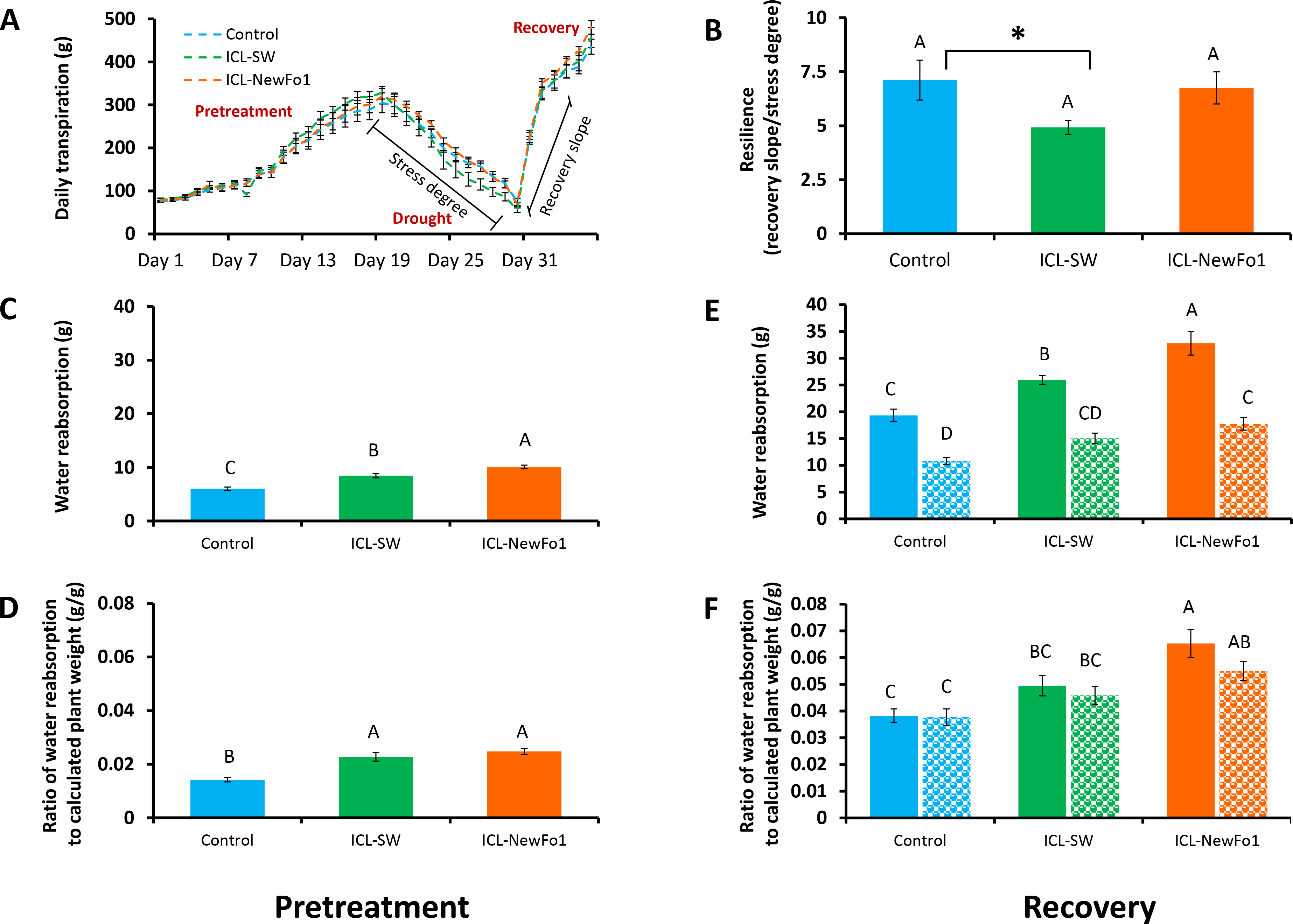
Effect of biostimulants on plant resilience during recovery. **(A)** Mean ± SE continuous total whole-plant daily transpiration of drought-treated plants during the entire experimental period of 36 days. Graph shows days from when stress degree and recovery slope were calculated for analysis. **(B)** Mean ± SE resilience measured as the ratio of the recovery slope (day 31–32) to stress degree (day 18–30). **(C)** Mean ± SE water reabsorption during pretreatment (day 11–14), and **(D)** its mean ± SE normalized to calculated plant weight. **(E)** Mean ± SE water reabsorption during recovery phase (day 33–36), and **(F)** its mean ± SE normalized to calculated plant weight. Blue bars – untreated (with biostimulants) control plants; green bars – ICL-SW-treated plants; orange bars – ICL-NewFo1-treated plants. Solid bars – well-irrigated conditions; stippled bars – drought conditions. Groups were compared using ANOVA by Tukey HSD test and Student’s *t*◻test. Different letters and asterisk above columns represent significant differences (*P* < 0.05). Each mean ± SE is from at least 8 plants per group.

### Biostimulant Effect on Fruit Number

As pepper plants are indeterminate, we decided to terminate the experiment shortly after recovery, despite the fact that full fruit weight potential had not been reached. Nevertheless, at this stage, fruit set in all groups was assumed to reflect the treatment, as seen in the distribution of the three different fruit sizes (small, medium and commercial) (Supplementary Figure 3). For the well-irrigated plants, 33% of the control fruit reached a commercial size, compared to only 19% of ICL-SW-treated and 14% of ICL-NewFo1-treated plants’ fruit. The total number of fruit was counted for all six groups and correlated to cumulative daily transpiration (Figure 6). ICL-SW significantly enhanced the total fruit number under ample irrigation (Student’s *t*-test); however, the ICL-SW-treated plants were significantly affected by the drought relative to their well-irrigated condition (Figure 6A). As similar results were observed for the transpiration rate of these treated plants (Figure 3E), we calculated the correlation between total fruit number and cumulative daily transpiration. The correlation for well-irrigated plants was slightly better (R^2^ = 0.5) than that for plants subjected to drought (Figure 6B).

**FIGURE 6.**
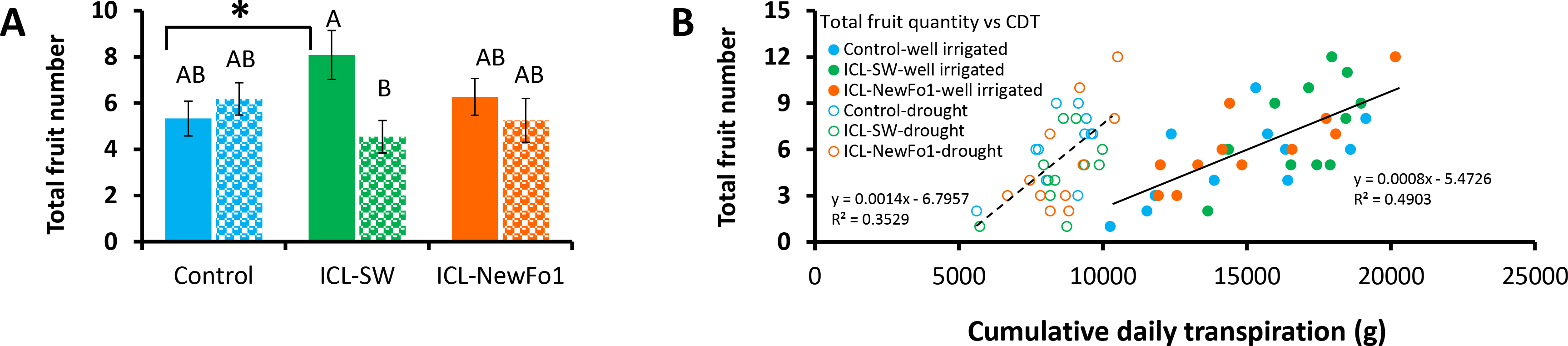
Effect of biostimulants on yield. **(A)** Mean ± SE total fruit number per plant. **(B)** Correlation between mean ± SE total fruit number and cumulative daily transpiration. Blue bars – untreated (with biostimulants) control plants; green bars – ICL-SW-treated plants; orange bars – ICL-NewFo1-treated plants. Solid bars – well-irrigated conditions; stippled bars – drought conditions. Groups were compared using ANOVA by Tukey HSD test and Student’s *t*◻test. Different letters and asterisk above columns represent significant differences (*P* < 0.05). Each mean ± SE is from at least 8 plants per group.

## DISCUSSION

### Advantages of the HFPS in Pre-Field Screening for Promising Candidates and Effective Treatments

Most high-throughput phenotyping facilities are based on remote sensing or imagers (Araus et al., 2018), and are expected to show improved temporal phenotypic resolution. However, their effective spatial resolution is still relatively limited to morphological and indirect physiological traits. In addition, measurements are not taken simultaneously; given the fact that the plant response to a dynamic environment is dynamic, simultaneous measurements are needed for comparative analyses. Thus, the selection of candidates and treatments for testing remains a challenge. High-throughput phenotyping platforms in greenhouses have the advantage of characterizing individual pot-grown plants without the constraints imposed by overlapping canopies from neighboring plants or variable climatic conditions that can hamper data-acquisition accuracy (Fernandez et al., 2017). Although an effective approach would be to screen biostimulants for their mode of action from “field to greenhouse”, the “greenhouse-to-field” approach is not only time- and cost-effective but also narrows the number of products to be tested later under field conditions (Rouphael et al., 2018). On the other hand, the accuracy of controlled growth environments in targeting genetically complex traits is questionable, as phenotypes from spaced pots and controlled conditions are poorly correlated with phenotypes in field environments, where plants compete with their neighbors (Nelissen et al., 2014; Poorter et al., 2016; Fernandez et al., 2017; Fischer et al., 2018; Rebetzke et al., 2018). We suggest that to better correlate a plant’s response to its environment, it is important to phenotype under conditions that are as similar as possible to those in the field. Thus, an efficient pre-field phenotype-screening experiment should offer the possibility to predict yield penalties in response to environmental adversity in the early stages of plant growth. Choice of the appropriate phenotyping method is one of the key components in pre-field screens (phenotyping) for complex traits under abiotic stress conditions (reviewed by Negin, 2017). This improves the chances of the selected candidates performing well under field conditions. The following principles, tested in this study, may contribute to this goal.

i. Conducting experiments under semi-controlled conditions that are typical of farmers’ growing facilities (see Figure 2A). The spaces between the pots were kept to a minimum to mimic commercial growth conditions.
ii. Using a truly randomized experimental design to mimic the biological variability, as well as the spatial and temporal variability in ambient conditions in the growth facility. Here, we used a randomized experimental design with one-to-one (1:1) plant– [sensors+controller] units which enabled running an independent feedback irrigation scenario for every plant (Figure 1, Supplementary Figure 1). Each controller was associated with a dual-valve system that allowed creating a specific combination of irrigation scenarios independently for each plant. Moreover, it overcame many of the experimental artifacts that could result from the “pot effect” (Gosa et al., 2018) by using controlled-deficit irrigation that reduced the irrigation levels every day to 80% of the previous day’s transpiration (for each plant separately and based on its individual performance), preventing rapid reduction in pot soil water content. This created a relatively homogeneous drought scenario for all plants (Figure 2).
iii. Conducting comparative and continuous measurements for all plants’ water-relations kinetics (direct physiological traits) in response to the three-phase scenario (control–drought–recovery). This experimental approach offered several advantages in interpreting the plant’s interactions with the environment as it compared each plant’s profile to its own profile in the different phases (Figure 3A) as well as to all other plants’ profiles in the experiment, simultaneously. Moreover, clarity of the stress conditions, providing the ability to repeat the exact stress scenario in other experiments, is also important when studying a desired stress-related trait. The trait in question might respond differently in plants showing different types of drought tolerance under different drought conditions (Negin and Moshelion, 2017). Therefore, for better resolution of the drought response in pot experiments, the severity and strategy of the drought stress must be well defined. To achieve a quantitative and cooperative response of the plants to a combination of biostimulants and drought treatment, we divided the experiment into three phases: before drought (pretreatment), during drought which was defined by the physiological drought point (θc), and recovery immediately after drought (resilience).
iv. High temporal and spatial resolution of the plant–environment interactions. The ultimate trait, yield, is a cumulative trait, measured at the end of the experiment and reflecting the sum of all genetic and ambient parameters affecting the plant throughout the season. This calls for high temporal resolution and continuous measurement of the dynamic plant–environment response. The high-capacity data acquisition (480 measurements per day) of the HFPS enabled tight measurements of the plant’s response to the ambient conditions, and also comparing plant performances at different time points during the day (i.e., different ambient conditions), where the differences between the treatments became significant (Figures 3 D,E).

In this study, we show that the HFPS might be an efficient diagnostic tool for a better understanding of pre-field plant x environment interactions by studying water-related physiological mechanisms under different phases of control–drought–recovery scenarios. In this pursuit, we used biostimulants as a test case due to their reported impact on the plant stress response (Van Oosten et al., 2017). Nevertheless, information on the influence of biostimulants on physiological mechanisms of action is relatively scarce. Moreover, the use of biostimulants, as with other biotic and abiotic screening studies, is highly complex, and thus identification and characterization of their activity is time-consuming and expensive, as it requires large-scale field experiments. The combination of the two types of treatment (biostimulants and drought) enabled us to evaluate the benefits of the HFPS in investigating the mechanistic effect of biostimulants in drought tolerance.

### Quantitative Comparison to Understand the Interactions between Key Physiological Traits and Their Trade-Offs

Both biostimulants increased plant transpiration rate under ample irrigation compared to control plants (Figure 3). However, while the impact of ICL-SW translated to productive mechanisms (faster growth rate and later, higher fruit number), the impact of ICL-NewFo1 translated to survivability mechanisms (lower transpiration rate under drought and faster recovery—i.e., better resilience). Interestingly, the sensitivity of the plants to drought in terms of the critical VWC drought point (θc), at which a further reduction in water content reduces transpiration, remained the same, possibly due to similar root sizes (Supplementary Figure 4). This is because under water-deficit conditions, when water becomes less available to the roots, plants with smaller roots will be limited more quickly (early θc) than plants with larger roots (reviewed by Gosa et al., 2018). Thus θc might be useful in predicting root phenotype. Nevertheless, as soon as the plants were exposed to drought, the two biostimulants induced different response patterns (beyond θc): ICL-NewFo1 treatment resulted in a gradual reduction in transpiration rate, reaching a minimum at a relatively lower VWC than the control and ICL-SW-treated plants (Figure 3C). Again, this type of behavior could explain the better survivability of the ICL-NewFo1-treated plants during the drought period.

In addition, the functional phenotyping approach revealed good correlations among key agronomical traits within the short study period. For example, our results revealed a high correlation between plant total dry weight and plant weight calculated by the system (Supplementary Figure 5). The fact that the system can calculate the plant biomass throughout the experiment is highly beneficial as it enables a direct measurement of the whole-plant biomass gain, in real time and in a non-destructive manner. In addition, key agronomic traits (such as grain yield) are linearly correlated to water consumption (WUE, reviewed by Gosa et al., 2018). Indeed, throughout the entire experiment, water taken up by the system (representing plant agronomic WUE, slope in Figure 4C) was almost identical to the fresh-weight WUE (taken on the first few days of the experiment; Figure 4E). Namely, ~0.003 g of plant dry weight per 1 mL of transpired water vs. ~0.035 g of plant fresh weight per 1 mL of transpired water, respectively, showing a ratio of 1:10, which is similar to the ratio between the fresh and dry shoot weight (Supplementary Figure 6). This trait is highly beneficial as it enables use of the fresh-weight WUE (determined on the first few days of the experiment), which is calculated for the entire growth period rather than the dry-weight WUE. Interestingly, these results also indicate that WUE is nearly constant throughout the growth period.

### Phenotyping Resilience

Resilience is one of the key stress-response traits. Nevertheless, the term “resilience” is being used more and more freely, and with popularity comes confusion; thus, it must assume its broadest definition. Resilience is commonly used to represent resistance, or recovery, or both (Hodgson et al., 2015). Plant stress resilience indicates plant survival and productivity after stress. In this study, we introduced two functional traits to quantify resilience: (i) transpiration recovery rate after stress (return of irrigation) and (ii) the plant’s ability to reabsorb water at night during recovery from drought. We found that while the biostimulants did not affect the transpiration recovery rate, they did increase the nighttime water reabsorption ability of both the well-irrigated and recovering plants, compared to the non-treated controls (Figures 5C,E). This phenomenon can be explained by the positive impact on the fresh biomass (Figure 4A), as normalizing the water reabsorption volume to the plant biomass still resulted in higher values of both biostimulant-treated plants compared to the non-treated control (Figures 5E,F). The difference between the water reabsorption of well-irrigated and recovering plants within the same group (i.e., control, ICL-SW or ICL-NewFo1) indicated drought-inflicted tissue damage, thus the night water reabsorption trait can be used as a tool to estimate tissue damage due to stress.

## CONCLUSION

A comparison of the effects of two biostimulants on drought tolerance using a HFPS revealed known and new relationships between physiological traits. The two studied biostimulants (ICL-SW and ICL-NewFo1) improved the overall transpiration and biomass gain compared to control plants. However, only ICL-SW improved fruit number (Figure 6A) under ample irrigation, which was significantly reduced when the plants were exposed to drought. This might be explained by the shift in resource allocation from the reproductive to non-reproductive or vegetative biomass, for survival. A schematic depiction of the behavior of plants treated with biostimulants is given in Figure 7. The behavior can be explained in terms of risk-taking and non-risk-taking behavior. Under optimal conditions, ICL-SW-treated plants (risk-taking) sustained a longer period of higher transpiration rate and thus a longer period of substantial CO_2_ assimilation, resulting in increased productivity (Figure 7A) compared to the ICL-NewFo1-treated plants. This behavior is advantageous only under well-irrigated conditions or during mild stress, but there is a risk of losing water faster during severe stress (Lin et◻al. 2007; Peng et◻al. 2007; McDowell et◻al. 2008; Sade et◻al. 2009; Moshelion et al., 2015). On the other hand, ICL-NewFo1-treated plants (non-risk-taking) maintain a moderate transpiration rate under optimal conditions, thus not contributing much to their productivity, but resulting in more gradual water loss under drought conditions, thereby reaching the minimal VWC (desiccation) later than the ICL-SW-treated plants, resulting in increased survivability. Thus there is a trade-off between productivity and survivability for the ICL-SW- and ICL-NewFo1-treated plants, respectively, as depicted in Figure 7B (Moshelion et al., 2015).

**FIGURE 7.**
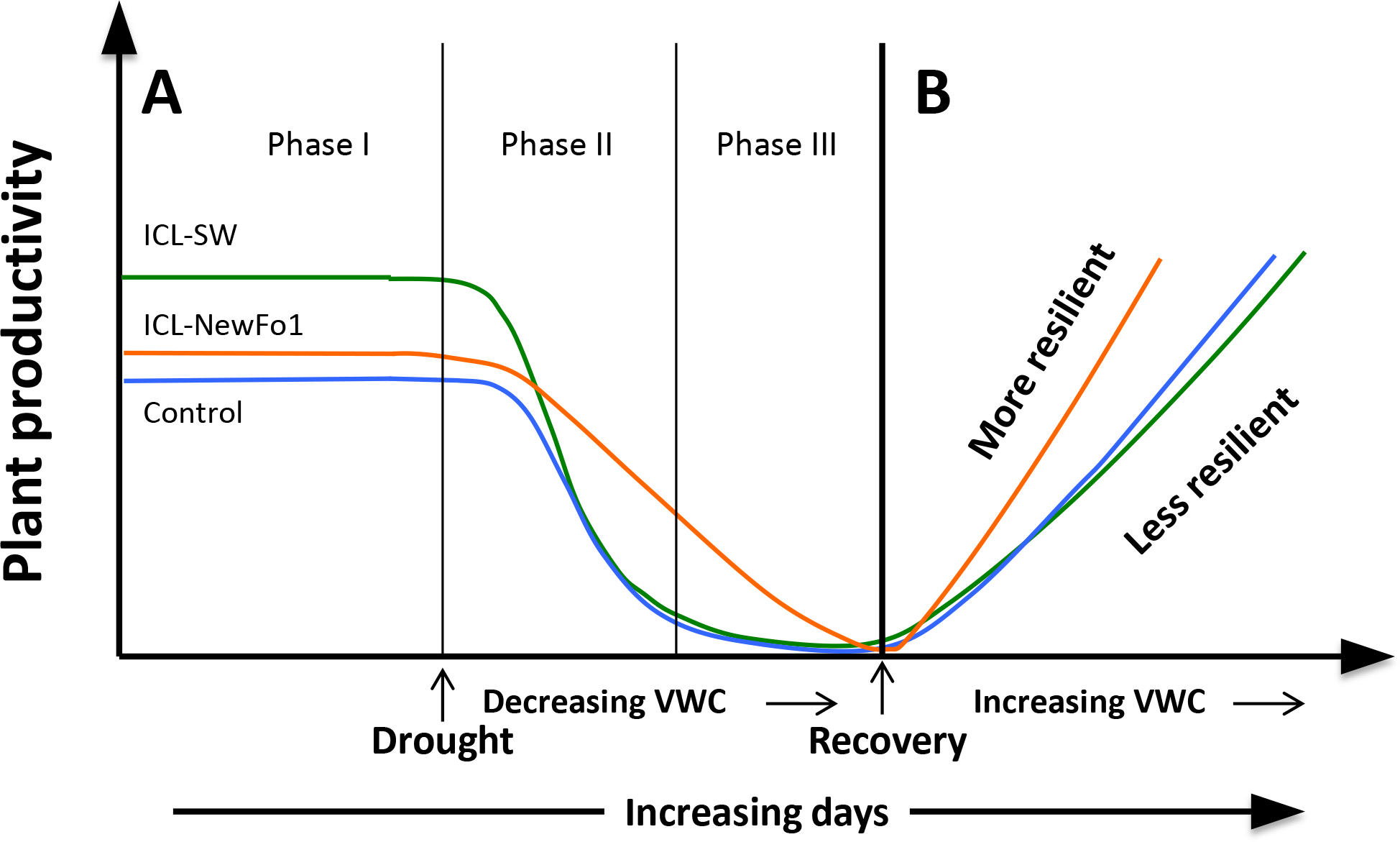
Schematic model of plant responses to biostimulants (ICL-NewFo1 – orange line, ICL-SW – green line, untreated – blue line) under drought and recovery (modified from Moshelion et al., 2015). **(A)** Plant productivity vs. intensity and duration of stress. Under conditions characterized by an ample water supply, ICL-SW-treated plants have a higher transpiration level than ICL-NewFo1-treated and control plants, and thus higher levels of productivity (e.g., photosynthesis) (Phase I). As mild water stress develops (Phase II), ICL-SW-treated and control plants reduce transpiration steeply with decreasing water availability, limiting productivity. In contrast, ICL-NewFo1-treated plants show a relatively gradual decrease in transpiration and productivity as a trade-off to the decline in leaf water potential and relative water content. Nevertheless, after the initial drought (Phase II), their productivity may still be higher than that of ICL-SW-treated and control plants which have already reached minimal productivity. As drought stress becomes more severe (Phase III), the transpiration values and productivity of ICL-NewFo1-treated plants continue to decline to their minimum. **(B)** Evaluation of recovery from drought is an important step in assessing drought resilience. It reveals the plant’s resistance to desiccation and ability to recover its pre-stress productivity, reflecting the extent of the damage caused by severe drought, such as cavitation or leaf/root loss. Both the ICL-SW-treated and control plants recover slowly compared to the ICL-NewFo1-treated plants. ICL-NewFo1 contributes to drought resistance by inducing more gradual water loss and resilience, thus contributing less to plant productivity and more to plant survivability. However, ICL-SW induces relatively faster water loss, and only increases productivity under optimal conditions while having no effect on the survivability of the plants under drought.

We suggest that these two different stimulation approaches should be implemented in different agricultural practices. Thus, the beneficial stimuli of ICL-SW may be implemented in controlled-irrigated crops, while the resilience impact of ICL-NewFo1 can be implemented for non-irrigated crops that are naturally subjected to the uncertainty of the environment. This survivability trait may also be very beneficial for annual crops (e.g., vines, turfs and silviculture) which need to overcome longer stress periods between seasons.

## AUTHOR CONTRIBUTIONS

RW and MM conceived the original research plan. AD, NH and IS performed the experiments. RB adapted the algorithms suggested by Halperin et al. (2017) to the calculations performed in this manuscript. AD, RB and MM analyzed the data. AD and MM wrote the manuscript. All authors were involved in reviewing and editing the manuscript.

## ACKNOWLEDGMENTS

AD and MM gratefully acknowledge the United States–Israel Binational Science Foundation (BSF) for the postdoctoral scholarship awarded to AD.

## References

Araus, J.L. and Cairns, J.E., 2014. Field high-throughput phenotyping: the new crop breeding frontier. Trends in plant science, 19(1), pp.52–61.

Araus, J.L., Kefauver, S.C., Zaman-Allah, M., Olsen, M.S. and Cairns, J.E., 2018. Translating high-throughput phenotyping into genetic gain. Trends in plant science.

Basak, A. (2008). “Biostimulators – definitions, classification and legislation,” in Monographs Series: Biostimulators in Modern Agriculture. General Aspects ed H. Gawrońska (Warsaw: Wieś Jutra), 7–17.

Bhat, J.A., Ali, S., Salgotra, R.K., Mir, Z.A., Dutta, S., Jadon, V., Tyagi, A., Mushtaq, M., Jain, N., Singh, P.K. and Singh, G.P., 2016. Genomic selection in the era of next generation sequencing for complex traits in plant breeding. Frontiers in genetics 7, p.221.

Brown, P. and Saa, S., 2015. Biostimulants in agriculture. Frontiers in plant science 6, p.671.

Bulgari, R., Cocetta, G., Trivellini, A., Vernieri, P., and Ferrante, A. (2015). Biostimulants and crop responses: a review. Biol. Agric. Hortic. 31, 1–17. doi: 10.1080/01448765.2014.964649

Collard, B.C. and Mackill, D.J., 2008. Marker-assisted selection: an approach for precision plant breeding in the twenty-first century. Philosophical Transactions of the Royal Society of London B: Biological Sciences 363(1491), pp.557–572.

Dalal, A., Attia, Z. and Moshelion, M., 2017. To Produce or to Survive: How Plastic Is Your Crop Stress Physiology?. Frontiers in plant science 8, p.2067.

Du Jardin, P. (2012). The Science of Plant Biostimulants - A Bibliographic Analysis, Ad hoc Study Report. Brussels: European Commission. Available online at: http://hdl.handle.net/2268/169257 (Accessed April 25, 2013).

du Jardin, P., 2015. Plant biostimulants: definition, concept, main categories and regulation. Scientia Horticulturae 196, pp.3–14.

Fernandez, M.G.S., Bao, Y., Tang, L. and Schnable, P.S., 2017. A high-throughput, field-based phenotyping technology for tall biomass crops. Plant physiology pp.pp-00707.

Fiorani, F. and Schurr, U., 2013. Future scenarios for plant phenotyping. Annual review of plant biology 64, pp.267–291.

Fischer, R.A., Byerlee, D. and Edmeades, G., 2014. Crop yields and global food security. ACIAR: Canberra, ACT pp.8–11.

Fischer, R.A. and Rebetzke, G.J., 2018. Indirect selection for potential yield in early-generation, spaced plantings of wheat and other small-grain cereals: a review. Crop and Pasture Science 69(5), pp.439–459.

Ghanem, M.E., Marrou, H. and Sinclair, T.R., 2015. Physiological phenotyping of plants for crop improvement. Trends in Plant Science 20(3), pp.139–144.

Gosa, S.C., Lupo, Y. and Moshelion, M., 2018. Quantitative and comparative analysis of whole-plant performance for functional physiological traits phenotyping: New tools to support pre-breeding and plant stress physiology studies. Plant Science.

Halperin, O., Gebremedhin, A., Wallach, R. and Moshelion, M., 2017. High[throughput physiological phenotyping and screening system for the characterization of plant–environment interactions. The Plant Journal 89(4), pp.839–850.

Hodgson, D., McDonald, J.L. and Hosken, D.J., 2015. What do you mean,‘resilient’?. Trends in ecology & evolution 30(9), pp.503–506.

Ikrina, M. A., and Kolbin, A. M. (2004). Regulators of Plant Growth and Development, Vol. 1, Stimulants. Moscow: Chimia.

Kumar, J., Pratap, A. and Kumar, S. eds., 2015. Phenomics in crop plants: trends, options and limitations (No. 8, p. 296). Springer India.

Li, L., Zhang, Q. and Huang, D., 2014. A review of imaging techniques for plant phenotyping. Sensors 14(11), pp.20078–20111.

Lin, W., Peng, Y., Li, G., Arora, R., Tang, Z., Su, W. and Cai, W., 2007. Isolation and functional characterization of PgTIP1, a hormone-autotrophic cells-specific tonoplast aquaporin in ginseng. Journal of experimental botany 58(5), pp.947–956.

McDowell, N., Pockman, W.T., Allen, C.D., Breshears, D.D., Cobb, N., Kolb, T., Plaut, J., Sperry, J., West, A., Williams, D.G. and Yepez, E.A., 2008. Mechanisms of plant survival and mortality during drought: why do some plants survive while others succumb to drought?. New phytologist 178(4), pp.719–739.

Miflin, B., 2000. Crop improvement in the 21st century. Journal of experimental botany 51(342), pp.1–8.

Moshelion, M. and Altman, A., 2015. Current challenges and future perspectives of plant and agricultural biotechnology. Trends in biotechnology 33(6), pp.337–342.

Moshelion, M., Halperin, O., Wallach, R., Oren, R.A.M. and Way, D.A., 2015. Role of aquaporins in determining transpiration and photosynthesis in water-stressed plants: crop water◻use efficiency, growth and yield. Plant, Cell & Environment 38(9), pp.1785–1793.

Negin, B. and Moshelion, M., 2017. The advantages of functional phenotyping in pre-field screening for drought-tolerant crops. Functional Plant Biology 44(1), pp.107–118.

Nelissen, H., Moloney, M. and Inzé, D., 2014. Translational research: from pot to plot. Plant biotechnology journal 12(3), pp.277–285.

Peng, Y., Lin, W., Cai, W. and Arora, R., 2007. Overexpression of a Panax ginseng tonoplast aquaporin alters salt tolerance, drought tolerance and cold acclimation ability in transgenic Arabidopsis plants. Planta 226(3), pp.729–740.

Petrozza, A., Santaniello, A., Summerer, S., Di Tommaso, G., Di Tommaso, D., Paparelli, E., Piaggesi, A., Perata, P. and Cellini, F., 2014. Physiological responses to Megafol^®^ treatments in tomato plants under drought stress: a phenomic and molecular approach. Scientia Horticulturae 174, pp.185–192.

Poorter, H., Fiorani, F., Pieruschka, R., Wojciechowski, T., Putten, W.H., Kleyer, M., Schurr, U. and Postma, J., 2016. Pampered inside, pestered outside? Differences and similarities between plants growing in controlled conditions and in the field. New Phytologist 212(4), pp.838–855.

Rahaman, M., Chen, D., Gillani, Z., Klukas, C. and Chen, M., 2015. Advanced phenotyping and phenotype data analysis for the study of plant growth and development. Frontiers in plant science 6, p.619.

Rebetzke, G.J., Jimenez-Berni, J., Fischer, R.A., Deery, D.M. and Smith, D.J., 2018. High-throughput phenotyping to enhance the use of crop genetic resources. Plant Science.

Rouphael, Y., Spíchal, L., Panzarová, K., Casa, R. and Colla, G., 2018. High-Throughput Plant Phenotyping for Developing Novel Biostimulants: From Lab to Field or From Field to Lab?. Frontiers in plant science 9.

Sade, N., Vinocur, B.J., Diber, A., Shatil, A., Ronen, G., Nissan, H., Wallach, R., Karchi, H. and Moshelion, M., 2009. Improving plant stress tolerance and yield production: is the tonoplast aquaporin SlTIP2; 2 a key to isohydric to anisohydric conversion?. New Phytologist 181(3), pp.651–661.

Salekdeh, G.H., Reynolds, M., Bennett, J. and Boyer, J., 2009. Conceptual framework for drought phenotyping during molecular breeding. Trends in plant science 14(9), pp.488–496.

Spindel, J., Begum, H., Akdemir, D., Virk, P., Collard, B., Redona, E., Atlin, G., Jannink, J.L. and McCouch, S.R., 2015. Genomic selection and association mapping in rice (Oryza sativa): effect of trait genetic architecture, training population composition, marker number and statistical model on accuracy of rice genomic selection in elite, tropical rice breeding lines. PLoS genetics 11(2), p.e1004982.

Tardieu, F., Cabrera-Bosquet, L., Pridmore, T. and Bennett, M., 2017. Plant phenomics, from sensors to knowledge. Current Biology 27(15), pp.R770–R783.

Van Oosten, M.J., Pepe, O., De Pascale, S., Silletti, S. and Maggio, A., 2017. The role of biostimulants and bioeffectors as alleviators of abiotic stress in crop plants. Chemical and Biological Technologies in Agriculture 4(1), p.5.

White, J.W., Andrade-Sanchez, P., Gore, M.A., Bronson, K.F., Coffelt, T.A., Conley, M.M., Feldmann, K.A., French, A.N., Heun, J.T., Hunsaker, D.J. and Jenks, M.A., 2012. Field-based phenomics for plant genetics research. Field Crops Research 133, pp.101–112.

Yakhin, O.I., Lubyanov, A.A., Yakhin, I.A. and Brown, P.H., 2017. Biostimulants in plant science: a global perspective. Frontiers in plant science 7, p.2049.

Yoo, C.Y., Pence, H.E., Hasegawa, P.M. and Mickelbart, M.V., 2009. Regulation of transpiration to improve crop water use. Critical Reviews in Plant Science 28(6), pp.410–431.

