## Supplementary Figure for "A High-Throughput Physiological Functional Phenotyping System for Time- and Cost-Effective Screening of Potential Biostimulants"

### Slide 1
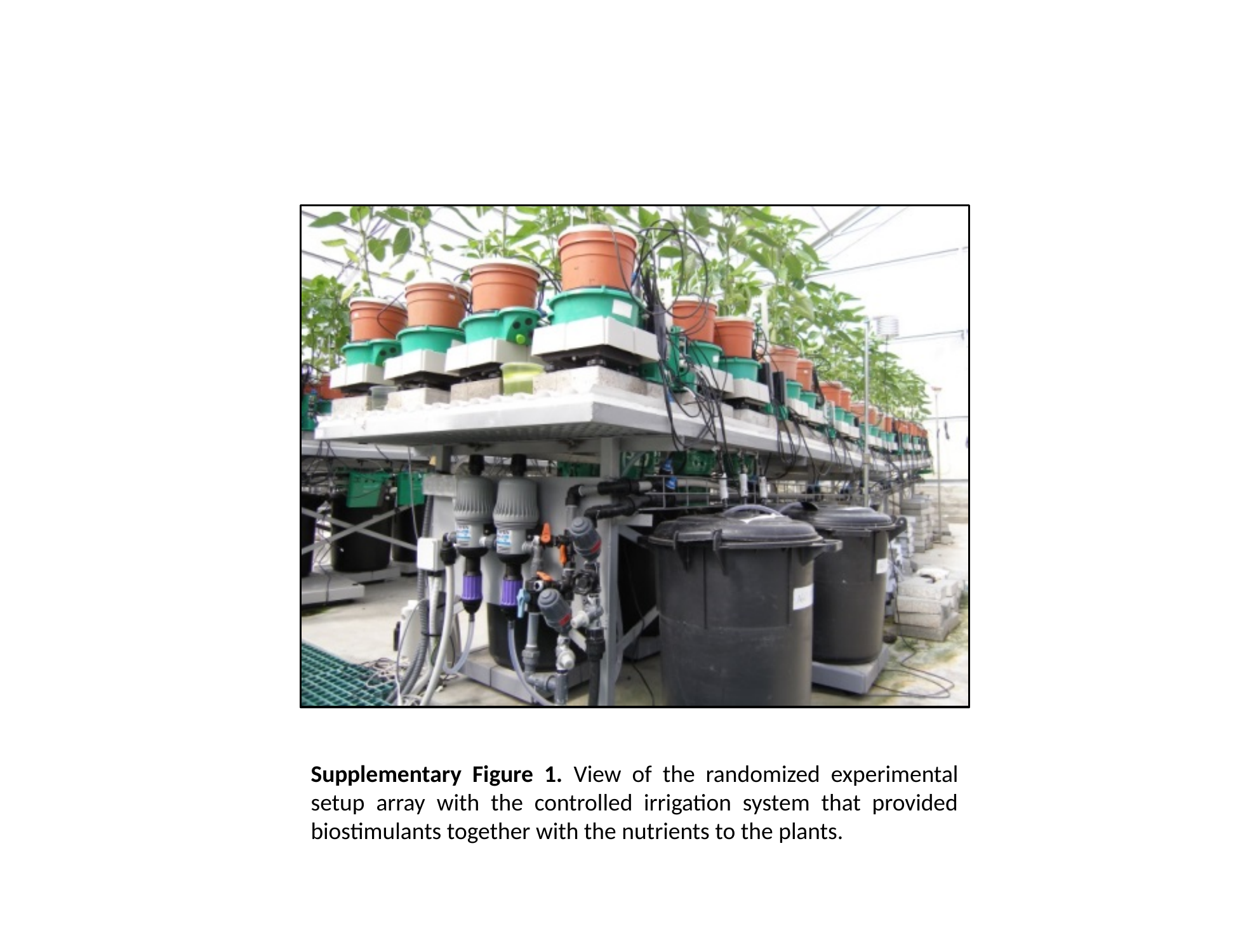

Supplementary Figure 1. View of the randomized experimental setup array with the controlled irrigation system that provided biostimulants together with the nutrients to the plants.

### Slide 2
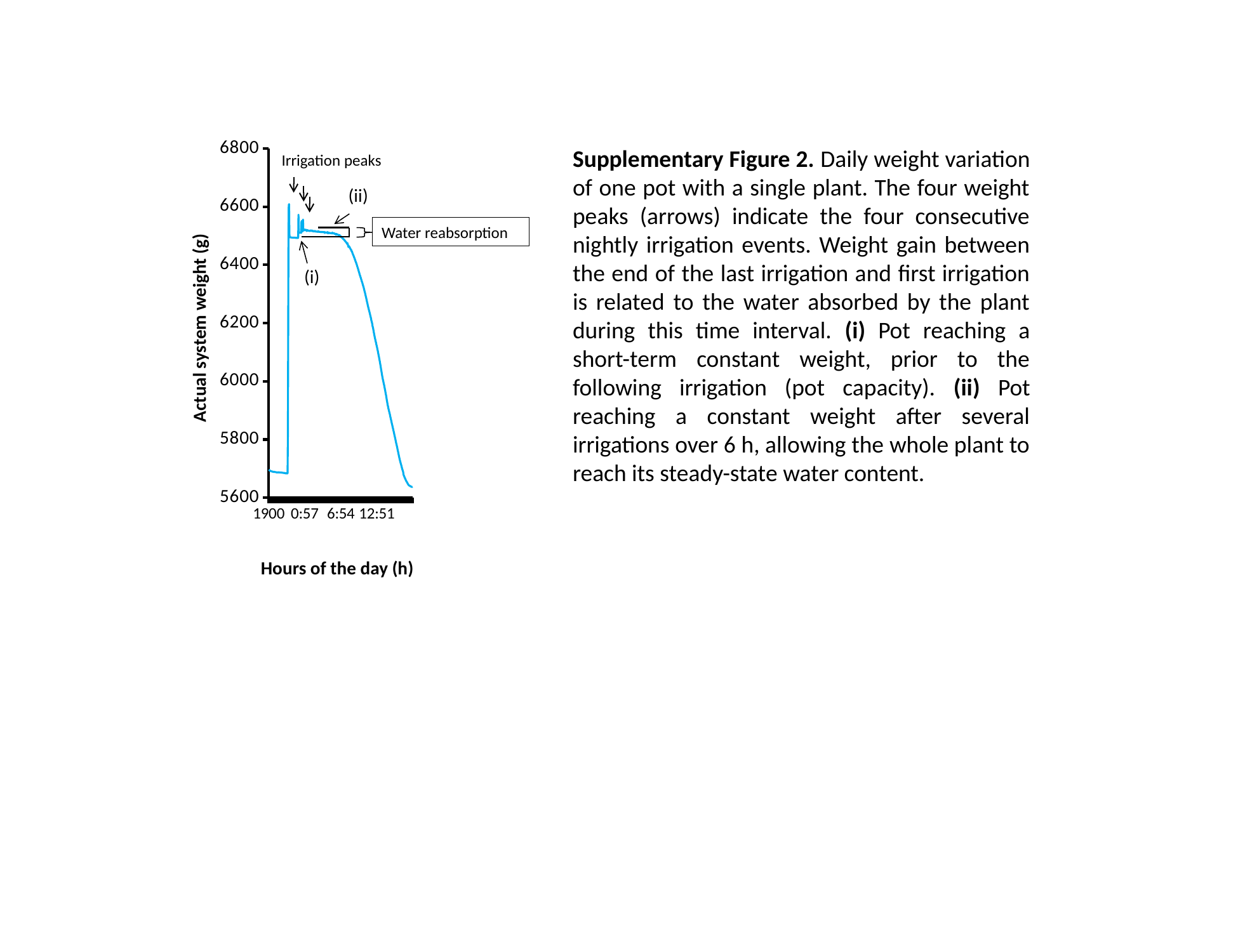

#### Chart
| Category | |
|---|---|
| 1900 | 5696.3 |
| 19:03 | 5695.22 |
| 19:06 | 5694.0 |
| 19:09 | 5692.91 |
| 19:12 | 5692.79 |
| 19:15 | 5692.41 |
| 19:18 | 5691.55 |
| 19:21 | 5690.88 |
| 19:24 | 5690.47 |
| 19:27 | 5690.48 |
| 19:30 | 5690.27 |
| 19:33 | 5689.75 |
| 19:36 | 5689.43 |
| 19:39 | 5689.3 |
| 19:42 | 5688.92 |
| 19:45 | 5688.49 |
| 19:48 | 5688.58 |
| 19:51 | 5688.59 |
| 19:54 | 5688.64 |
| 19:57 | 5688.57 |
| 20:00 | 5687.85 |
| 20:03 | 5687.87 |
| 20:06 | 5687.23 |
| 20:09 | 5687.75 |
| 20:12 | 5687.42 |
| 20:15 | 5687.03 |
| 20:18 | 5686.96 |
| 20:21 | 5686.69 |
| 20:24 | 5686.71 |
| 20:27 | 5686.49 |
| 20:30 | 5687.07 |
| 20:33 | 5687.14 |
| 20:36 | 5686.79 |
| 20:39 | 5686.98 |
| 20:42 | 5687.36 |
| 20:45 | 5686.46 |
| 20:48 | 5686.56 |
| 20:51 | 5686.41 |
| 20:54 | 5686.31 |
| 20:57 | 5686.28 |
| 21:00 | 5686.24 |
| 21:03 | 5686.51 |
| 21:06 | 5685.98 |
| 21:09 | 5685.81 |
| 21:12 | 5685.77 |
| 21:15 | 5685.28 |
| 21:18 | 5685.74 |
| 21:21 | 5685.21 |
| 21:24 | 5685.31 |
| 21:27 | 5684.57 |
| 21:30 | 5684.91 |
| 21:33 | 5684.84 |
| 21:36 | 5684.83 |
| 21:39 | 5685.04 |
| 21:42 | 5683.77 |
| 21:45 | 5683.68 |
| 21:48 | 5683.93 |
| 21:51 | 5683.71 |
| 21:54 | 5683.64 |
| 21:57 | 5683.56 |
| 22:00 | 5683.11 |
| 22:03 | 5683.32 |
| 22:06 | 5683.38 |
| 22:09 | 5800.74 |
| 22:12 | 6126.55 |
| 22:15 | 6505.19 |
| 22:18 | 6604.99 |
| 22:21 | 6608.94 |
| 22:24 | 6505.36 |
| 22:27 | 6495.62 |
| 22:30 | 6494.61 |
| 22:33 | 6494.56 |
| 22:36 | 6494.51 |
| 22:39 | 6494.24 |
| 22:42 | 6494.28 |
| 22:45 | 6494.19 |
| 22:48 | 6493.7 |
| 22:51 | 6494.2 |
| 22:54 | 6493.65 |
| 22:57 | 6493.75 |
| 2300 | 6493.44 |
| 23:03 | 6493.5 |
| 23:06 | 6493.08 |
| 23:09 | 6493.38 |
| 23:12 | 6493.53 |
| 23:15 | 6493.52 |
| 23:18 | 6493.62 |
| 23:21 | 6493.33 |
| 23:24 | 6493.02 |
| 23:27 | 6493.06 |
| 23:30 | 6492.44 |
| 23:33 | 6492.59 |
| 23:36 | 6492.95 |
| 23:39 | 6492.59 |
| 23:42 | 6492.37 |
| 23:45 | 6492.76 |
| 23:48 | 6493.22 |
| 23:51 | 6492.39 |
| 23:54 | 6572.33 |
| 23:57 | 6514.0 |
| 0:00 | 6510.84 |
| 0:03 | 6510.42 |
| 0:06 | 6510.56 |
| 0:09 | 6510.32 |
| 0:12 | 6510.93 |
| 0:15 | 6510.77 |
| 0:18 | 6510.26 |
| 0:21 | 6509.99 |
| 0:24 | 6550.51 |
| 0:27 | 6517.61 |
| 0:30 | 6514.9 |
| 0:33 | 6515.22 |
| 0:36 | 6515.66 |
| 0:39 | 6555.62 |
| 0:42 | 6520.46 |
| 0:45 | 6518.92 |
| 0:48 | 6519.14 |
| 0:51 | 6518.68 |
| 0:54 | 6519.07 |
| 0:57 | 6521.22 |
| 1:00 | 6519.51 |
| 1:03 | 6519.61 |
| 1:06 | 6520.6 |
| 1:09 | 6521.04 |
| 1:12 | 6519.77 |
| 1:15 | 6516.22 |
| 1:18 | 6517.89 |
| 1:21 | 6517.4 |
| 1:24 | 6519.64 |
| 1:27 | 6517.23 |
| 1:30 | 6517.49 |
| 1:33 | 6517.71 |
| 1:36 | 6518.0 |
| 1:39 | 6517.11 |
| 1:42 | 6517.02 |
| 1:45 | 6516.07 |
| 1:48 | 6517.51 |
| 1:51 | 6516.6 |
| 1:54 | 6517.29 |
| 1:57 | 6518.7 |
| 2:00 | 6516.57 |
| 2:03 | 6515.79 |
| 2:06 | 6516.8 |
| 2:09 | 6518.04 |
| 2:12 | 6516.13 |
| 2:15 | 6517.41 |
| 2:18 | 6515.07 |
| 2:21 | 6516.41 |
| 2:24 | 6514.77 |
| 2:27 | 6515.36 |
| 2:30 | 6515.12 |
| 2:33 | 6515.98 |
| 2:36 | 6516.74 |
| 2:39 | 6514.64 |
| 2:42 | 6514.26 |
| 2:45 | 6515.11 |
| 2:48 | 6515.14 |
| 2:51 | 6513.61 |
| 2:54 | 6514.32 |
| 2:57 | 6514.01 |
| 0300 | 6514.51 |
| 3:03 | 6513.79 |
| 3:06 | 6513.73 |
| 3:09 | 6515.77 |
| 3:12 | 6514.77 |
| 3:15 | 6513.88 |
| 3:18 | 6513.46 |
| 3:21 | 6513.52 |
| 3:24 | 6513.62 |
| 3:27 | 6514.27 |
| 3:30 | 6513.22 |
| 3:33 | 6513.16 |
| 3:36 | 6511.91 |
| 3:39 | 6512.58 |
| 3:42 | 6513.24 |
| 3:45 | 6514.05 |
| 3:48 | 6513.0 |
| 3:51 | 6512.43 |
| 3:54 | 6512.3 |
| 3:57 | 6513.08 |
| 4:00 | 6512.7 |
| 4:03 | 6512.43 |
| 4:06 | 6511.76 |
| 4:09 | 6511.71 |
| 4:12 | 6513.07 |
| 4:15 | 6509.63 |
| 4:18 | 6512.18 |
| 4:21 | 6511.0 |
| 4:24 | 6511.46 |
| 4:27 | 6511.67 |
| 4:30 | 6511.31 |
| 4:33 | 6511.04 |
| 4:36 | 6511.15 |
| 4:39 | 6509.5 |
| 4:42 | 6512.91 |
| 4:45 | 6511.14 |
| 4:48 | 6508.29 |
| 4:51 | 6510.23 |
| 4:54 | 6510.58 |
| 4:57 | 6510.66 |
| 5:00 | 6509.85 |
| 5:03 | 6509.38 |
| 5:06 | 6509.43 |
| 5:09 | 6509.38 |
| 5:12 | 6508.29 |
| 5:15 | 6509.48 |
| 5:18 | 6508.39 |
| 5:21 | 6509.16 |
| 5:24 | 6508.8 |
| 5:27 | 6507.7 |
| 5:30 | 6510.99 |
| 5:33 | 6509.15 |
| 5:36 | 6508.57 |
| 5:39 | 6509.0 |
| 5:42 | 6508.82 |
| 5:45 | 6507.72 |
| 5:48 | 6509.13 |
| 5:51 | 6506.95 |
| 5:54 | 6507.68 |
| 5:57 | 6506.23 |
| 6:00 | 6506.73 |
| 6:03 | 6506.46 |
| 6:06 | 6503.36 |
| 6:09 | 6506.35 |
| 6:12 | 6505.75 |
| 6:15 | 6505.24 |
| 6:18 | 6505.32 |
| 6:21 | 6504.36 |
| 6:24 | 6503.88 |
| 6:27 | 6503.45 |
| 6:30 | 6502.19 |
| 6:33 | 6502.19 |
| 6:36 | 6501.77 |
| 6:39 | 6502.03 |
| 6:42 | 6502.55 |
| 6:45 | 6499.18 |
| 6:48 | 6499.27 |
| 6:51 | 6496.1 |
| 6:54 | 6497.98 |
| 6:57 | 6497.12 |
| 0700 | 6495.72 |
| 7:03 | 6496.12 |
| 7:06 | 6492.83 |
| 7:09 | 6493.36 |
| 7:12 | 6491.91 |
| 7:15 | 6489.62 |
| 7:18 | 6489.04 |
| 7:21 | 6488.27 |
| 7:24 | 6487.7 |
| 7:27 | 6485.95 |
| 7:30 | 6484.91 |
| 7:33 | 6483.61 |
| 7:36 | 6482.18 |
| 7:39 | 6480.32 |
| 7:42 | 6480.12 |
| 7:45 | 6478.87 |
| 7:48 | 6477.17 |
| 7:51 | 6475.6 |
| 7:54 | 6473.87 |
| 7:57 | 6471.91 |
| 8:00 | 6474.89 |
| 8:03 | 6469.33 |
| 8:06 | 6468.01 |
| 8:09 | 6461.73 |
| 8:12 | 6464.98 |
| 8:15 | 6464.1 |
| 8:18 | 6462.74 |
| 8:21 | 6461.16 |
| 8:24 | 6458.21 |
| 8:27 | 6456.89 |
| 8:30 | 6454.93 |
| 8:33 | 6452.63 |
| 8:36 | 6452.17 |
| 8:39 | 6448.45 |
| 8:42 | 6446.64 |
| 8:45 | 6445.82 |
| 8:48 | 6442.1 |
| 8:51 | 6439.07 |
| 8:54 | 6436.63 |
| 8:57 | 6433.67 |
| 9:00 | 6431.57 |
| 9:03 | 6429.26 |
| 9:06 | 6426.06 |
| 9:09 | 6423.67 |
| 9:12 | 6420.83 |
| 9:15 | 6417.65 |
| 9:18 | 6414.33 |
| 9:21 | 6411.3 |
| 9:24 | 6408.63 |
| 9:27 | 6406.14 |
| 9:30 | 6402.57 |
| 9:33 | 6399.75 |
| 9:36 | 6396.01 |
| 9:39 | 6392.94 |
| 9:42 | 6389.78 |
| 9:45 | 6386.21 |
| 9:48 | 6383.22 |
| 9:51 | 6378.93 |
| 9:54 | 6375.61 |
| 9:57 | 6372.17 |
| 10:00 | 6367.39 |
| 10:03 | 6364.63 |
| 10:06 | 6361.8 |
| 10:09 | 6358.13 |
| 10:12 | 6355.42 |
| 10:15 | 6351.75 |
| 10:18 | 6348.64 |
| 10:21 | 6344.51 |
| 10:24 | 6341.78 |
| 10:27 | 6338.83 |
| 10:30 | 6334.76 |
| 10:33 | 6331.49 |
| 10:36 | 6327.4 |
| 10:39 | 6324.2 |
| 10:42 | 6319.88 |
| 10:45 | 6315.53 |
| 10:48 | 6311.32 |
| 10:51 | 6307.48 |
| 10:54 | 6303.49 |
| 10:57 | 6298.87 |
| 1100 | 6294.43 |
| 11:03 | 6290.02 |
| 11:06 | 6285.86 |
| 11:09 | 6280.81 |
| 11:12 | 6276.29 |
| 11:15 | 6271.37 |
| 11:18 | 6266.83 |
| 11:21 | 6261.47 |
| 11:24 | 6257.0 |
| 11:27 | 6253.24 |
| 11:30 | 6248.86 |
| 11:33 | 6244.62 |
| 11:36 | 6240.51 |
| 11:39 | 6236.89 |
| 11:42 | 6232.4 |
| 11:45 | 6228.11 |
| 11:48 | 6223.76 |
| 11:51 | 6219.59 |
| 11:54 | 6215.44 |
| 11:57 | 6208.63 |
| 12:00 | 6205.52 |
| 12:03 | 6200.77 |
| 12:06 | 6195.92 |
| 12:09 | 6188.37 |
| 12:12 | 6185.87 |
| 12:15 | 6180.1 |
| 12:18 | 6174.84 |
| 12:21 | 6168.54 |
| 12:24 | 6162.83 |
| 12:27 | 6157.43 |
| 12:30 | 6152.54 |
| 12:33 | 6147.6 |
| 12:36 | 6143.06 |
| 12:39 | 6137.86 |
| 12:42 | 6133.61 |
| 12:45 | 6128.44 |
| 12:48 | 6123.89 |
| 12:51 | 6119.4 |
| 12:54 | 6115.2 |
| 12:57 | 6109.53 |
| 13:00 | 6105.06 |
| 13:03 | 6099.82 |
| 13:06 | 6094.29 |
| 13:09 | 6088.78 |
| 13:12 | 6083.83 |
| 13:15 | 6078.28 |
| 13:18 | 6072.76 |
| 13:21 | 6066.69 |
| 13:24 | 6060.37 |
| 13:27 | 6054.67 |
| 13:30 | 6048.33 |
| 13:33 | 6041.28 |
| 13:36 | 6035.75 |
| 13:39 | 6029.28 |
| 13:42 | 6022.75 |
| 13:45 | 6017.04 |
| 13:48 | 6011.77 |
| 13:51 | 6006.08 |
| 13:54 | 6001.8 |
| 13:57 | 5996.46 |
| 14:00 | 5991.79 |
| 14:03 | 5986.88 |
| 14:06 | 5981.31 |
| 14:09 | 5976.05 |
| 14:12 | 5970.85 |
| 14:15 | 5965.62 |
| 14:18 | 5959.19 |
| 14:21 | 5954.4 |
| 14:24 | 5948.3 |
| 14:27 | 5942.18 |
| 14:30 | 5935.25 |
| 14:33 | 5929.83 |
| 14:36 | 5923.46 |
| 14:39 | 5918.12 |
| 14:42 | 5912.78 |
| 14:45 | 5907.6 |
| 14:48 | 5903.08 |
| 14:51 | 5898.8 |
| 14:54 | 5894.74 |
| 14:57 | 5890.27 |
| 1500 | 5886.55 |
| 15:03 | 5881.12 |
| 15:06 | 5876.22 |
| 15:09 | 5872.22 |
| 15:12 | 5867.28 |
| 15:15 | 5862.91 |
| 15:18 | 5857.93 |
| 15:21 | 5853.25 |
| 15:24 | 5848.65 |
| 15:27 | 5843.78 |
| 15:30 | 5839.24 |
| 15:33 | 5834.85 |
| 15:36 | 5830.17 |
| 15:39 | 5825.58 |
| 15:42 | 5821.2 |
| 15:45 | 5816.12 |
| 15:48 | 5811.36 |
| 15:51 | 5806.8 |
| 15:54 | 5802.19 |
| 15:57 | 5797.52 |
| 16:00 | 5792.17 |
| 16:03 | 5787.38 |
| 16:06 | 5782.83 |
| 16:09 | 5778.09 |
| 16:12 | 5773.67 |
| 16:15 | 5768.27 |
| 16:18 | 5763.3 |
| 16:21 | 5758.53 |
| 16:24 | 5754.0 |
| 16:27 | 5748.35 |
| 16:30 | 5743.46 |
| 16:33 | 5738.79 |
| 16:36 | 5734.31 |
| 16:39 | 5729.71 |
| 16:42 | 5725.56 |
| 16:45 | 5721.36 |
| 16:48 | 5717.36 |
| 16:51 | 5713.95 |
| 16:54 | 5710.13 |
| 16:57 | 5706.17 |
| 17:00 | 5701.82 |
| 17:03 | 5698.15 |
| 17:06 | 5694.88 |
| 17:09 | 5691.56 |
| 17:12 | 5688.38 |
| 17:15 | 5679.8 |
| 17:18 | 5676.89 |
| 17:21 | 5674.39 |
| 17:24 | 5671.93 |
| 17:27 | 5668.7 |
| 17:30 | 5666.8 |
| 17:33 | 5664.26 |
| 17:36 | 5662.19 |
| 17:39 | 5659.61 |
| 17:42 | 5657.9 |
| 17:45 | 5655.35 |
| 17:48 | 5654.16 |
| 17:51 | 5652.43 |
| 17:54 | 5650.33 |
| 17:57 | 5648.15 |
| 18:00 | 5646.62 |
| 18:03 | 5645.17 |
| 18:06 | 5643.21 |
| 18:09 | 5642.56 |
| 18:12 | 5641.33 |
| 18:15 | 5641.26 |
| 18:18 | 5640.45 |
| 18:21 | 5640.01 |
| 18:24 | 5639.35 |
| 18:27 | 5638.18 |
| 18:30 | 5637.96 |
| 18:33 | 5637.68 |
| 18:36 | 5637.41 |
| 18:39 | 5636.99 |
| 18:42 | 5636.51 |
| 18:45 | 5637.2 |Supplementary Figure 2. Daily weight variation of one pot with a single plant. The four weight peaks (arrows) indicate the four consecutive nightly irrigation events. Weight gain between the end of the last irrigation and first irrigation is related to the water absorbed by the plant during this time interval. (i) Pot reaching a short-term constant weight, prior to the following irrigation (pot capacity). (ii) Pot reaching a constant weight after several irrigations over 6 h, allowing the whole plant to reach its steady-state water content.
Irrigation peaks
(ii)
Water reabsorption
(i)
Actual system weight (g)
Hours of the day (h)

### Slide 3
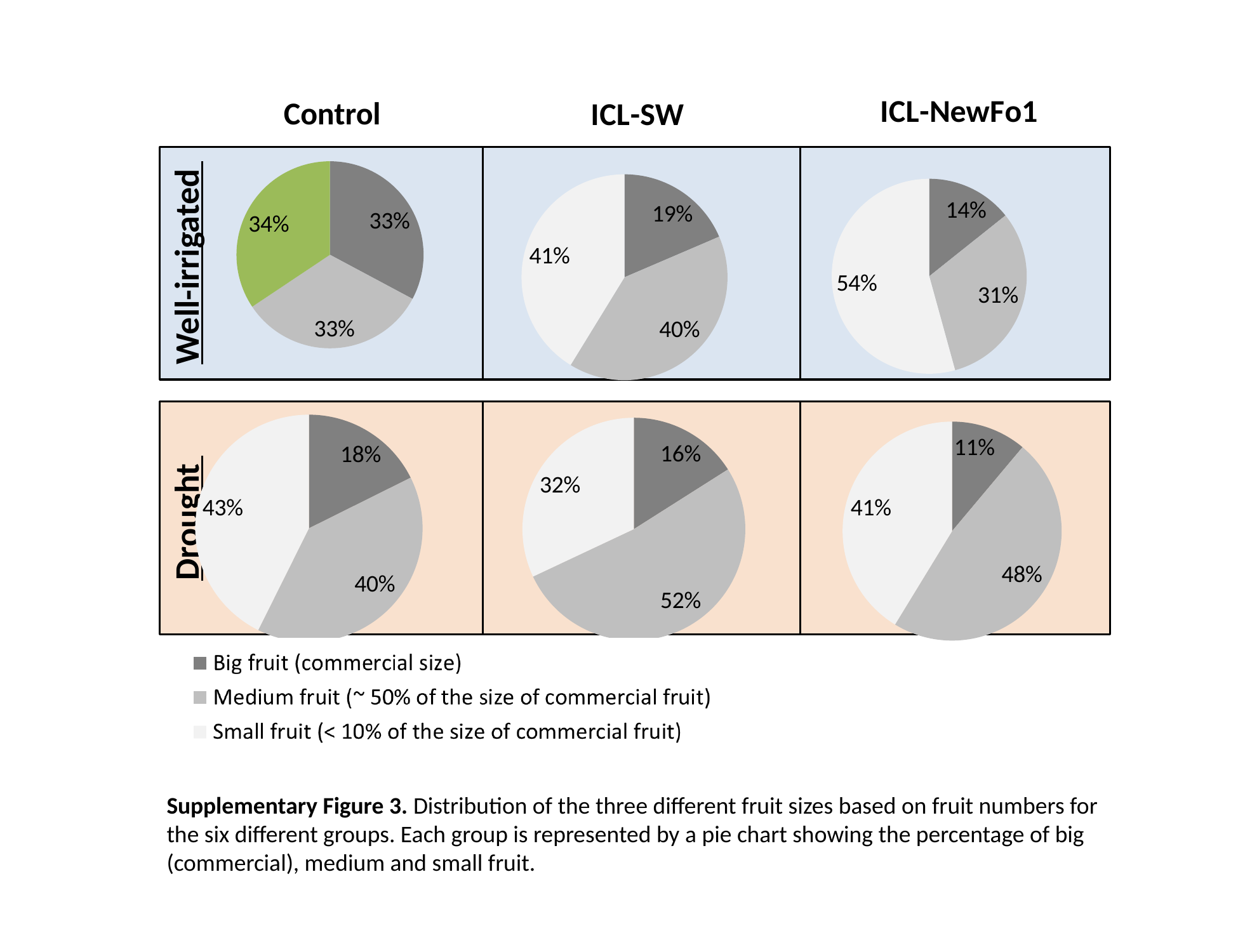

#### Chart: ICL-SW
| Category | |
|---|---|
| Big Fruit (Comercial size) | 1.5 |
| Medium Fruit (~ 50% of the size of commercial fruit) | 3.25 |
| Small Fruit (Less than 25% of the size of commercial fruit) | 3.3333333333333335 |
#### Chart: ICL-NewFo1
| Category | |
|---|---|
| Big Fruit (Comercial size) | 0.8333333333333334 |
| Medium Fruit (~ 50% of the size of commercial fruit) | 1.8333333333333333 |
| Small Fruit (Less than 25% of the size of commercial fruit) | 3.1666666666666665 |
#### Chart: Control
| Category | |
|---|---|
| Big Fruit (Comercial size) | 1.75 |
| Medium Fruit (~ 50% of the size of commercial fruit) | 1.75 |
| Small Fruit (Less than 25% of the size of commercial fruit) | 1.8333333333333333 |
Well-irrigated
#### Chart
| Category | |
|---|---|
| Big Fruit | 1.0909090909090908 |
| Medium Fruit | 2.4545454545454546 |
| Small Fruit | 2.6363636363636362 |
#### Chart
| Category | |
|---|---|
| Big Fruit (Comercial size) | 0.5833333333333334 |
| Medium Fruit (~ 50% of the size of commercial fruit) | 2.5 |
| Small Fruit (Less than 25% of the size of commercial fruit) | 2.1666666666666665 |
#### Chart
| Category | |
|---|---|
| Big Fruit (Comercial size) | 0.7272727272727273 |
| Medium Fruit (~ 50% of the size of commercial fruit) | 2.3636363636363638 |
| Small Fruit (Less than 25% of the size of commercial fruit) | 1.4545454545454546 |
Drought
Supplementary Figure 3. Distribution of the three different fruit sizes based on fruit numbers for the six different groups. Each group is represented by a pie chart showing the percentage of big (commercial), medium and small fruit.

### Slide 4
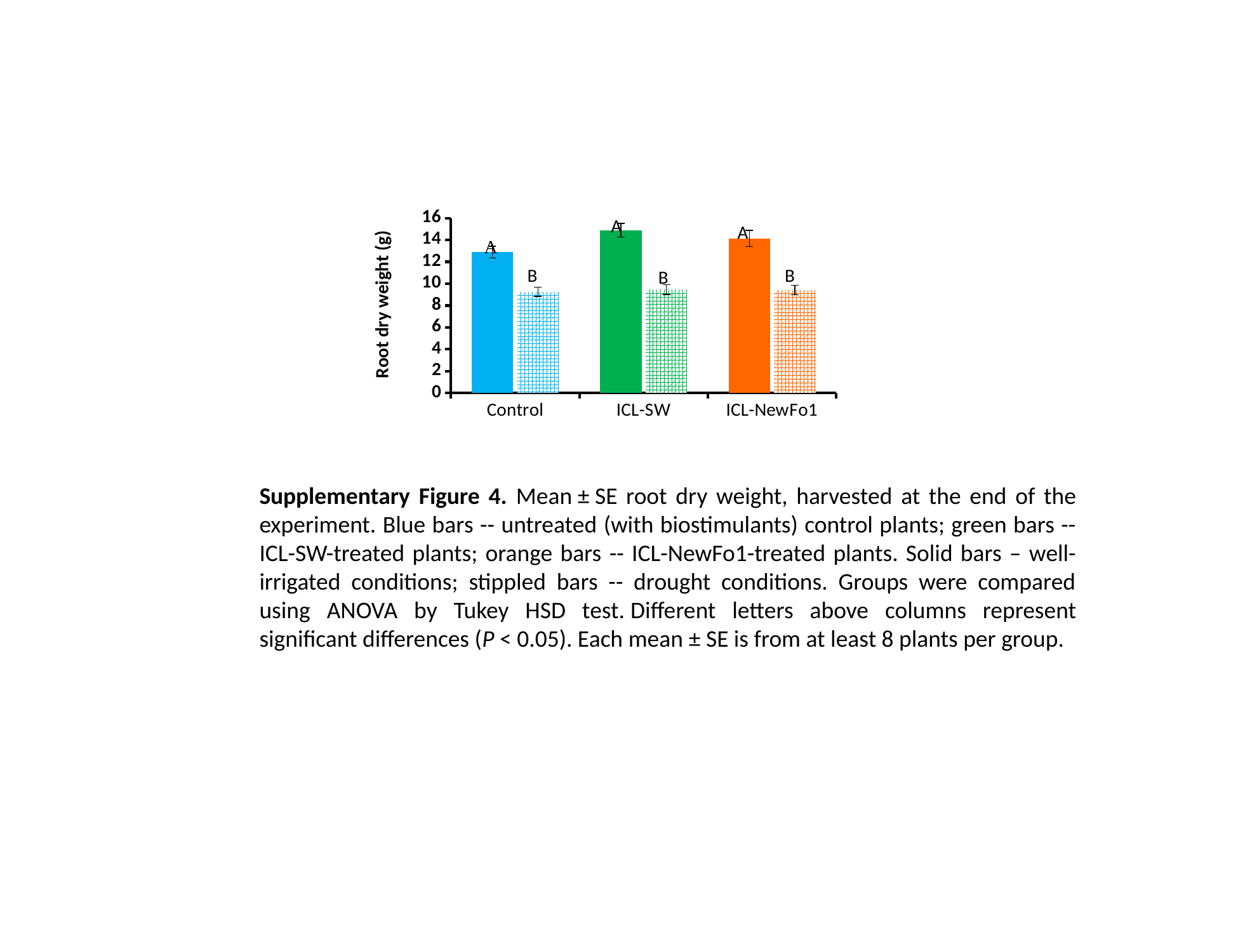

#### Chart
| Category | None | Drought |
|---|---|---|
| Control | 12.883333333 | 9.2636363636 |
| ICL-SW | 14.883333333 | 9.4545454545 |
| ICL-NewFo1 | 14.145454545 | 9.425 |Root dry weight (g)
Supplementary Figure 4. Mean ± SE root dry weight, harvested at the end of the experiment. Blue bars -- untreated (with biostimulants) control plants; green bars -- ICL-SW-treated plants; orange bars -- ICL-NewFo1-treated plants. Solid bars – well-irrigated conditions; stippled bars -- drought conditions. Groups were compared using ANOVA by Tukey HSD test. Different letters above columns represent significant differences (P < 0.05). Each mean ± SE is from at least 8 plants per group.

### Slide 5
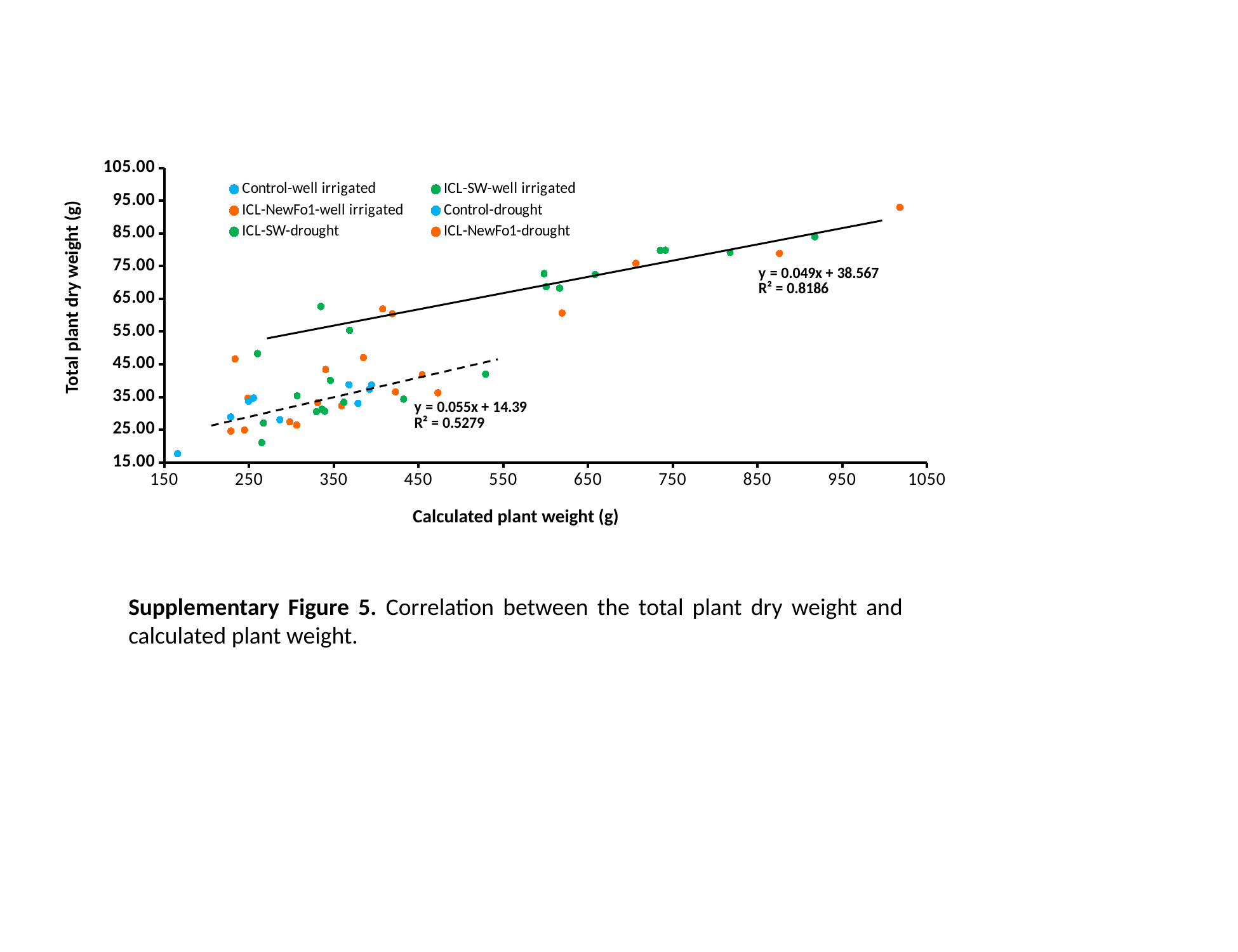

#### Chart
| Category | | | | | | |
|---|---|---|---|---|---|---|Total plant dry weight (g)
Calculated plant weight (g)
Supplementary Figure 5. Correlation between the total plant dry weight and calculated plant weight.

### Slide 6
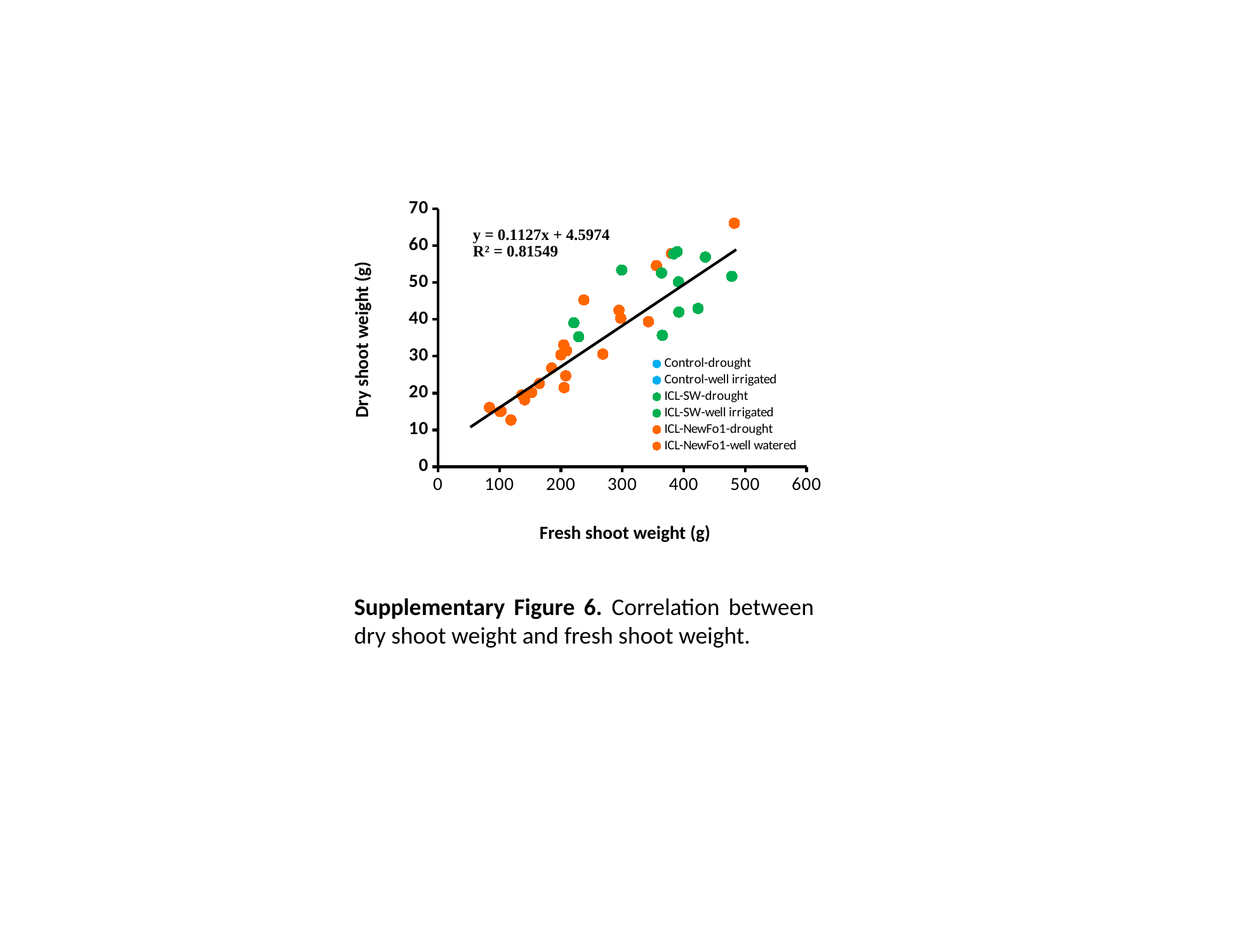

#### Chart
| Category | | | | | | |
|---|---|---|---|---|---|---|Dry shoot weight (g)
Fresh shoot weight (g)
Supplementary Figure 6. Correlation between dry shoot weight and fresh shoot weight.
