## Supplementary material for "A High-Throughput Physiological Functional Phenotyping System for Time- and Cost-Effective Screening of Potential Biostimulants": Table 1

**TABLE 1. Nutrient composition of irrigation solution before 20% dilution.**

| **Final concentration (mM)** | **Final concentration (ppm)** | **Mineral** |
| --- | --- | --- |
| 2.3 | 195.8 | NaNO_3_ (N) |
| 0.000969 | 209 | H_3_PO_4_ (P) |
| 2.685 | 271.4 | KNO_3_ (K) |
| 0.623 | 75 | MgSO_4_ (Mg) |
| 0.0025 | 0.748 | ZnSO_4_ (Zn) |
| 0.00198 | 0.496 | CuSO_4_ (CU) |
| 0.00081 | 0.131 | MoO_3_ (Mo) |
| 0.0154 | 3.441 | MnSO_4_ (Mn) |
| 0.00078 | 0.3 | Borax (B) |
| 0.0204 | 8.66 | C_10_H_2_FeN_2_NaO_8_ (Fe) |

Citric acid was used to reduce the pH of the stock solution (concentrated 100X from the values mentioned in the table) to 1.5. The pH of the final irrigation solution from the dripper (after dilution with tap water) varied between 6.5 and 7.
